# ErbB inhibition rescues nigral dopamine neuron hyperactivity and repetitive behaviors in a mouse model of fragile X syndrome

**DOI:** 10.1101/2024.02.23.581801

**Authors:** Sebastian Luca D’Addario, Eleonora Rosina, Mariangela Massaro Cenere, Claudia Bagni, Nicola Biagio Mercuri, Ada Ledonne

**Affiliations:** Department of Experimental Neuroscience, Santa Lucia Foundation IRCCS, Rome, Italy; Department of Biomedicine and Prevention, University of Rome Tor Vergata, Rome, Italy; Department of Fundamental Neurosciences, University of Lausanne, Lausanne, Switzerland; Neurology Unit, Department of Systems Medicine, University of Rome Tor Vergata, Rome, Italy; Pharmacology Unit, Department of Systems Medicine, University of Rome Tor Vergata, Rome, Italy

**Keywords:** ErbB signaling, neuregulins, dopamine, substantia nigra, mGluR1, hyperexcitability, *Fmr1* KO mice, self-grooming, marble burying, autism spectrum disorders

## Abstract

Repetitive behaviors are core symptoms of autism spectrum disorders (ASD) and fragile X syndrome (FXS), the prevalent genetic cause of intellectual disability and autism. The nigrostriatal dopamine (DA) circuit rules movement and habit formation; therefore, its dysregulation stands as a leading substrate for repetitive behaviors. However, beyond indirect evidence, specific assessment of nigral DA neuron activity in ASD and FXS models is lacking. Here, we show that hyperactivity of substantia nigra pars compacta (SNpc) DA neurons is an early feature of FXS. The underlying mechanisms rely on mGluR1 and ErbB receptors. Up-regulation of ErbB4 and ErbB2 in nigral DA neurons drives neuronal hyperactivity and repetitive behaviors of the FXS mouse, simultaneously rescued by ErbB inhibition. In conclusion, beyond providing the first evidence of dysregulation of the SNpc DA nucleus in FXS, we identify novel targets - ErbB receptors - whose inhibition proficiently attenuates repetitive behaviors, thus opening an avenue toward innovative therapies for ASD and FXS.

## Introduction

Repetitive behaviors, such as stereotyped movements, repetitive objects handling, and self-injurious behaviors, are core diagnostic signs of autism spectrum disorders (ASD), also overt in fragile X syndrome (FXS), the main genetic cause of autism and intellectual disability caused by *Fmr1* silencing and loss of Fragile X Messenger Ribonucleoprotein (FMRP)^1,2,3^. The nigrostriatal dopamine (DA) circuit, arising from DA neurons of the substantia nigra pars compacta (SNpc), is key to movement control and the creation of habits and sequential behaviors^4,5^; thus, its dysregulation is posited as a prominent substrate of abnormal movements and restricted routines, emerging as compulsive and stereotyped behaviors in patients with ASD and FXS. Earlier clues about the role of the SNpc DA nucleus in repetitive behaviors stand up from evidence that pharmacological and genetic manipulations increasing striatal DA transmission in rodents promote stereotyped movements by striatal D1 activation^6,7,8,9,10,11^. The latest demonstration that optogenetic activation of SNpc DA neurons triggers self-grooming (a repetitive behavior) in mice^12^ and self-grooming is attenuated by optogenetic inhibition of the SNpc-to-ventromedial striatum circuit^13^ has strengthened the idea that hyperactivation of the nigrostriatal DA circuit is instrumental for repetitive behaviors. However, so far, precise evidence proving altered activity of nigral DA neuron in ASD or FXS models is absent, keeping the involvement of the SNpc DA nucleus in abnormal repetitive behaviors still rather theoretical^14,15,16,17^.

Striatal dysfunctions have been recently reported in the *Fmr1* KO mouse, a validated model for FXS^18,19^. Few other studies described histological and neurochemical DA alterations in the striatum of this FXS model^17^: evidence is partially divergent, as histological analyses of the nigrostriatal DA circuit in adult *Fmr1* KO mice found increased branching of striatal tyrosine hydroxylase positive (TH+) terminals^20^ or loss of SNpc DA neurons^21^, while evaluation of striatal DA dynamics reported bidirectional age-dependent changes in this FXS model, with higher DA turnover at one month^22^, increased DA tissue levels at two months^23^, and a reduction of basal levels^24^ and electrically-evoked DA release^25^ in older mice. Due to these disjointed data and, more importantly, shortage of direct investigation of nigral DA neuron activity in ASD and FXS models, current knowledge about the role of the nigrostriatal DA circuit in ASD and FXS is inadequate. Furthermore, since evidence of striatal DA alterations in FXS models is mostly restricted to adulthood, the earlier DA-dependent mechanisms driving pediatric FXS symptoms are entirely unknown. In the present study, we directly assess whether the nigrostriatal DA circuit contributes to the etiological processes of FXS by analyzing the activity of nigral DA neurons in adolescent *Fmr1* KO mice. We show that hyperactivity of nigral DA neurons is an early signature of FXS, and the underlying mechanism relies on the interaction between metabotropic glutamate receptor 1 (mGluR1) and ErbB tyrosine kinases, receptors for the neurotrophic and differentiation factors known as neuregulins (NRGs). Lastly, we investigate whether pharmacological inhibition of ErbB signaling represents a valuable approach to rescue nigral DA neuron dysfunctions and repetitive behaviors in the FXS model.

## Results

### Hyperactivity of nigral dopamine neurons in a FXS model

To evaluate whether SNpc DA neuron activity is altered in FXS, we performed electrophysiological recordings in acute midbrain slices of male adolescent *Fmr1* KO and WT mice (**Fig. 1a**). First, we analyzed spontaneous firing activity in the patch clamp cell-attached configuration and observed that nigral DA neurons in the FXS mouse spontaneously fired action potentials (APs) at higher frequencies than WT (**Fig. 1b**). Analysis of the coefficient of variation of the interspike intervals (CV-ISIs) revealed a similar temporal distribution of spikes (**Fig. 1c**), supporting maintenance of pacemaker regularity despite increased firing rate. Nigral DA neuron hyperactivity was overt in the FXS mouse at postnatal days (PND) 21-23 **(Supplementary Fig. 1)**, and was also evident in the whole-cell mode (**Fig. 1d**). SNpc DA neurons from the FXS mouse also showed modifications in passive membrane properties, precisely increased cell capacitance (C_m_), but no changes in membrane resistance (R_m_) (**Fig. 1e**). To verify variations in neuronal excitability, we measured APs evoked by depolarizing currents, revealing that SNpc DA neurons from F*mr1* KO mice fired more APs than WT (**Fig. 1f**). Collectively, these data demonstrate early dysregulation of the nigrostriatal DA circuit in FXS, identifying SNpc DA neuron hyperactivity as a novel feature of FXS.

**Figure 1.**
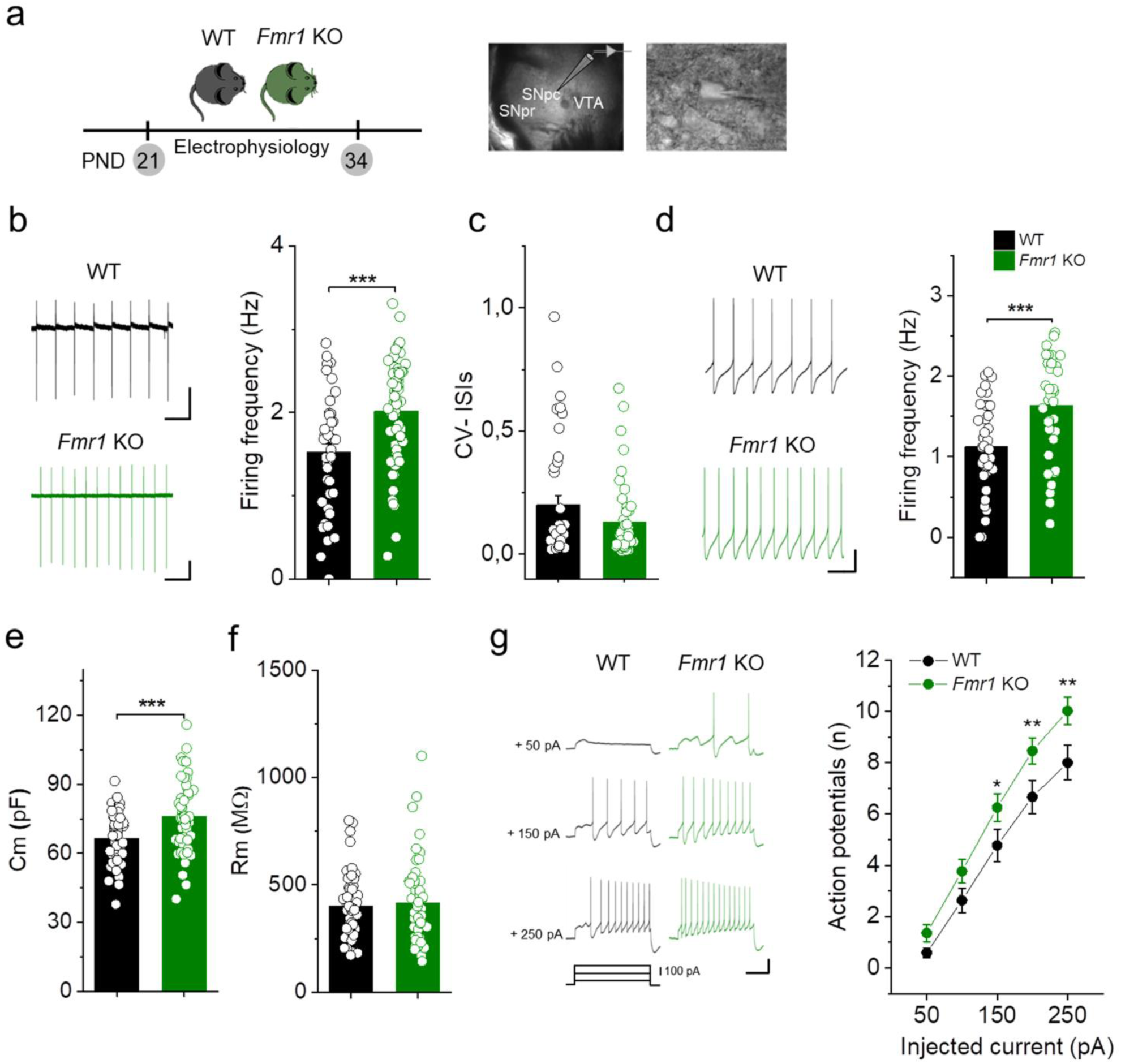
Hyperactivity of nigral dopamine neurons in a FXS model. **a)** Left, schematic of *ex vivo* electrophysiological recordings of nigral dopamine (DA) neurons in adolescent mice. Middle, image of an horizontal slice displaying the site of the recordings, the substantia nigra pars compacta (SNpc). Right, image of a SNpc DA neuron during the patch-clamp procedure. **b)** Left, example firing traces in cell-attached mode. Right, plot of mean firing frequency showing increased firing rate of SNpc DA neurons from *Fmr1* KO (n = 59 cells/10 mice) compared with WT mice (n = 41 cells/8 mice); P = 3.913 × 10^-4^, t = -3.671 and DF = 98, two-tailed unpaired *t*-test. Scale bar: 20 pA/1 s. **c)** Plot of coefficient of variation of interspike intervals (CV-ISIs) revealing similar temporal distribution of spikes between genotypes. WT (n = 40 cells/8 mice) and *Fmr1* KO (n = 48 cells/10 mice), P = 0.211, U = 1110, Z = 1.252, Mann-Whitney test. **d)** Left, example firing trace in whole-cell mode; Right, quantification of firing frequencies of WT (n = 36 cells/8 mice) and *Fmr1* KO mice (n = 38 cells/10 mice), P = 3.037 X10^-4^, Z = -3.531, U = 334, Mann-Whitney test. Scale bar: 20 mV/1s. **e)** Plot of membrane capacitance (C_m_) and **(f)** membrane resistance (R_m_) of nigral DA neurons from WT (n = 50 cells/8 mice) and *Fmr1* KO mice (n = 53 cells/10 mice). C_m_: P = 4.87 × 10^-4^, t = -3.604, DF = 101, two-tailed unpaired *t*-test; R_m_: P=0.711, two-tailed unpaired *t*-test. Scale bar: 20 mV/1s. **g)** Left, example traces of action potentials (APs) evoked by current injections (holding voltage V_H_ = -60 mV) in nigral DA neurons; Right, APs number at each current step. WT (n = 27 cells/7 mice) and *Fmr1* KO mice (n = 39 cells/9 mice). RM two-way ANOVA followed by Tukey’s test. Genotype: P = 0.047, injected current P=1.76 × 10^-50^, genotype X injected current P = 0.095. P = 0.023 at 150 pA, P=0.003 at 200 pA; P=0.001 at 250 pA. Scale bar: 25 mV/0.5 s. All data are presented as mean ± SEM.

### Abnormal mGluR1 function drives nigral DA neuron hyperactivity in the FXS mouse

The spontaneous firing activity of SNpc DA neurons is tightly controlled by several intrinsic and extrinsic factors that allow repetitive AP generation and pauses, resulting in tonic pacemaker activity in a narrow low-frequency range in *ex vivo* preparations^26^. In addition to intrinsic ion channels, afferent excitatory (E) and inhibitory (I) synaptic inputs can shape the firing rate. Since E/I imbalance is a common synaptic alteration underlying abnormal neuronal activity and circuitry connectivity in FXS^3,27^, we analyzed E/I ratio in nigral DA neurons by recording from the same neuron either spontaneous excitatory-(sEPSC, V_H_ = -70 mV) or inhibitory synaptic currents (sIPSC, V_H_ = +10 mV). We found no differences in frequency and amplitude of sEPSCs and sIPSCs **(Supplementary Fig. 2),** indicating normal E/I balance of fast synaptic transmission in the FXS model. These findings exclude that changes in glutamate or GABA release are major determinants of nigral DA neuron hyperactivity in the FXS mouse.

Group I metabotropic glutamate receptors (mGluRI) - mGluR1 and mGluR5 - control essential brain functions, influencing neuronal activity, neurotransmission, and synaptic plasticity in several brain areas^28,29^. While abnormal mGluR5 function is a recognized signature of FXS, the contribution of mGluR1 to the etiological processes of FXS is overlooked. As mGluR1 activation depolarizes SNpc DA neurons and enhances pacemaker and bursting firing^30,31,32,33^, we investigated whether mGluR1 hyperfunction is the functional substrate for nigral DA neuron hyperactivity in the FXS model. To test this, we first measured the depolarizing currents induced by the mGluR1/5 agonist DHPG (10 μM) (I_DHPG_), revealing that I_DHPG_ is potentiated in the nigral DA neurons of the FXS mouse (**Fig. 2a**). I_DHPG_ analyses in the presence of selective antagonists demonstrated that I_DHPG_ enhancement relies exclusively on mGluR1; a pretreatment with the mGluR1 antagonist CPCCOEt (100 µM), but not with the mGluR5 antagonist MPEP (10 µM), occluded I_DHPG_ differences between genotypes (**Fig. 2a**). Next, we evaluated if an abnormal mGluR1 function drives hyperactivity and hyperexcitability of nigral DA neurons in the FXS model. Blunting mGluR1 activity normalized spontaneous firing frequency (**Fig. 2b**) and excitability (**Fig. 2c**) of SNpc DA neurons in the FXS mouse. These findings reveal an unrecognized contribution of mGluR1 to the neurobiological mechanisms of FXS, showing that mGluR1 hyperfunction triggers early hyperactivation of the nigrostriatal DA circuit.

**Figure 2.**
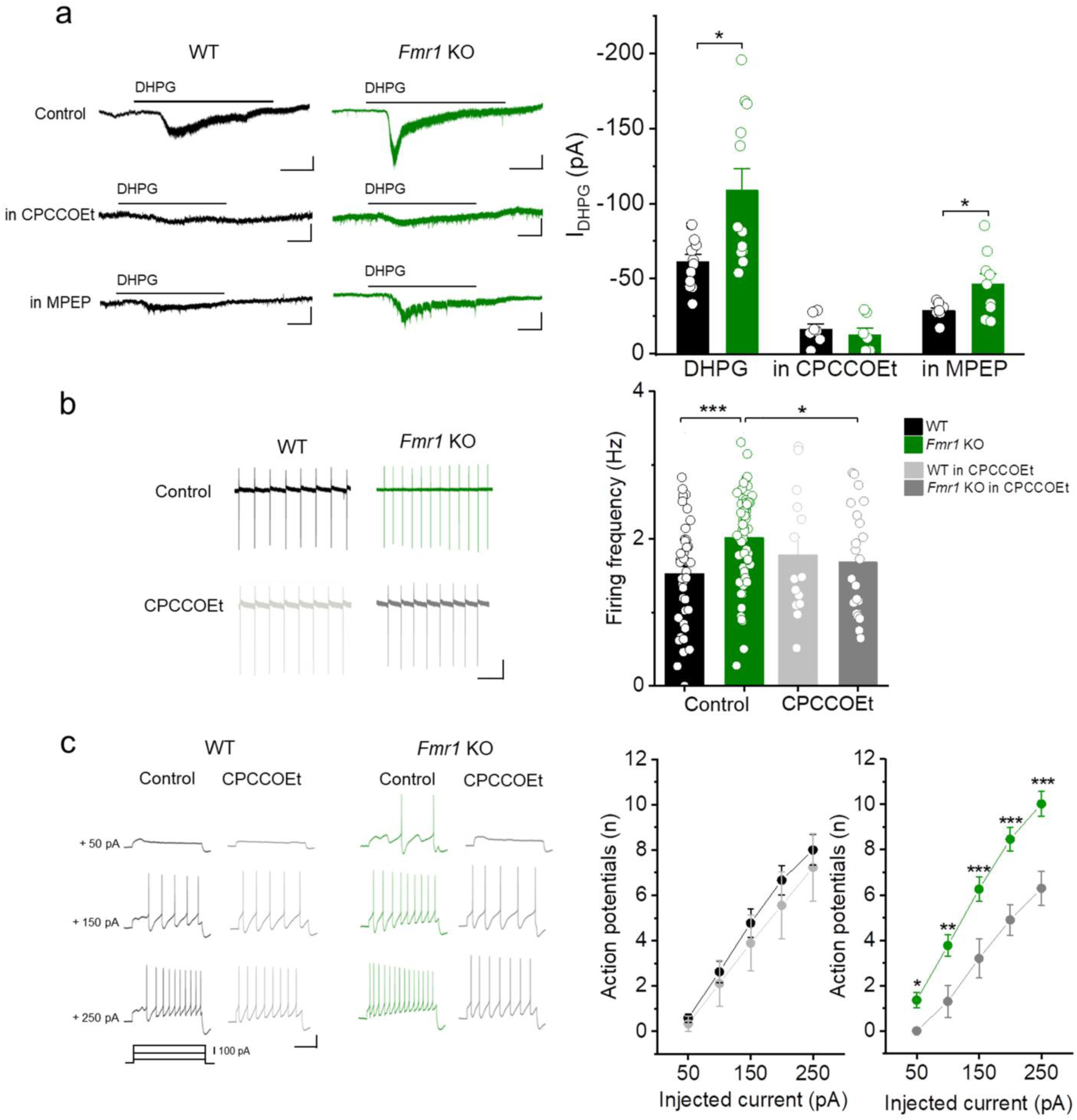
Exacerbated mGluR1 function drives nigral DA neuron hyperactivity in the FXS model. **a)** Left, example traces of currents activated by the mGluR1/5 agonist DHPG (10 µM) (I_DHPG_) in SNpc DA neurons in control and during mGluR1 or mGluR5 inhibition with CPCCOEt (100 µM) or MPEP (10 µM), respectively. Right, quantification of I_DHPG_ amplitudes showing increased I_DHPG_ in *Fmr1* KO mice (n = 12 cells/4 mice) compared with WT (n = 12 cells/3 mice). P = 0.023, U = 112, Mann-Whitney test. I_DHPG_ potentiation in the FXS mouse depends selectively on mGluR1. I_DHPG_ in CPCCOEt: WT (n = 7 cells/3 mice) and *Fmr1* KO (n = 7 cells/3 mice), P = 0.56, t = -0.597, and DF = 11, two-tailed unpaired *t*-test. I_DHPG_ in MPEP: WT (n = 8 cells/3 mice) and *Fmr1* KO (n = 9 cells/4 mice), P = 0.041, t = 2.228, and DF = 15, two-tailed unpaired *t*-test. Scale bar: 20 pA/ 0.5 s. **b)** Example spontaneous firing traces (left) and quantification of firing rates (right) showing that CPCCOEt attenuates SNpc DA neuron hyperactivity in the FXS mouse. WT-control (n = 41 cells/8 mice), *Fmr1* KO-control (n = 59 cells/10 mice), WT-CPCCOEt (n = 14 cells/5 mice), and *Fmr1* KO-CPCCOEt (n = 21 cells/5 mice). Two-way ANOVA followed by Fisher’s test. Genotype x drug interaction P = 0.026. P = 6.571 × 10^-4^ for WT-control vs *Fmr1* KO-control, P = 0.047 for *Fmr1* KO-control vs *Fmr1* KO-CPCCOEt. Scale bar: 20 pA/1s. **c)** Example traces (left) and plots of evoked APs (right) showing that inhibiting mGluR1 reduces SNpc DA neuron hyperexcitability in the FXS mouse. RM two-way ANOVA followed by Tukey’s test. CPCCOEt effect in WT: WT-control (n = 27 cells/7 mice) and WT-CPCCOEt (n = 9 cells/5 mice). Drug P = 0.635, injected current P = 1.331 × 10^-11^, drug X current interaction P = 0.304. CPCCOEt effect in *Fmr1* KO: *Fmr1* KO-control (n = 39 cells/10 mice) and *Fmr1* KO-CPCCOEt (n = 10 cells/5 mice). Drug P = 0.002, injected current P = 7.039 × 10^-17^, drug X current interaction P = 0.0277. P = 0.045 at 50 pA, P = 0.001 at 100 pA, P = 1.063 × 10^-4^ at 150 pA, P = 2.230 × 10^-5^ at 200 pA, P = 8.657 × 10^-6^ at 250 pA. Scale bar: 25 mV/0.5 s. All data are presented as mean ± SEM.

### Molecular determinants of mGluR1 hyperfunction

To assess whether mGluR1 hyperfunction in nigral DA neurons of the FXS mouse is associated with increased expression of mGluR1, we analyzed mGluR1 levels using the Western blot technique in homogenates of SNpc microdissected from 25-day-old adolescent mice (**Fig. 3a**). We found higher levels of nigral mGluR1 in the FXS mouse (**Fig. 3b**). Interestingly, such mGluR1 up-regulation appears confined to SNpc, since mGluR1 levels are unchanged in the adjacent DA nucleus, the ventral tegmental area (VTA) **(Supplementary Fig. 3)**.

**Figure 3.**
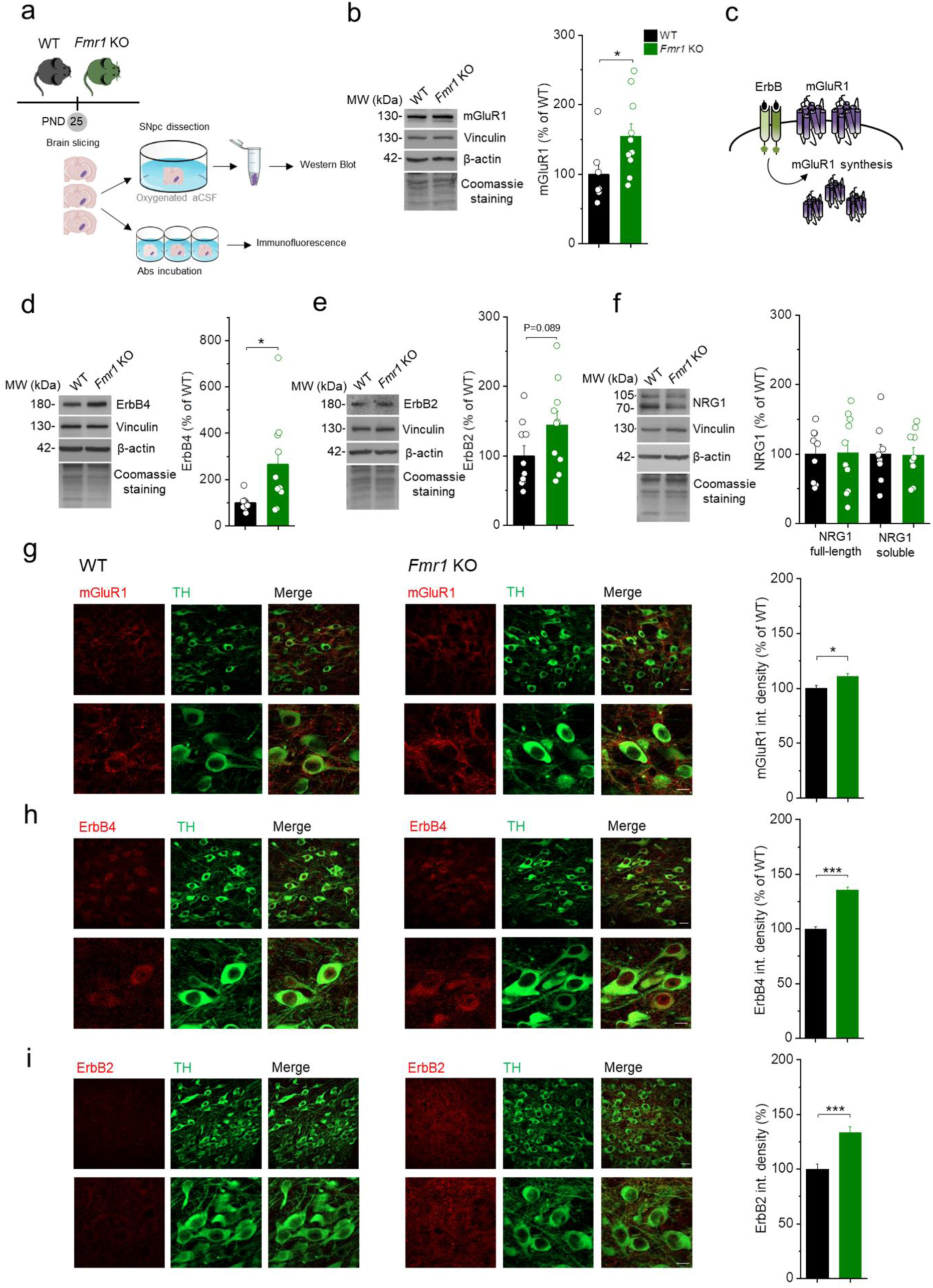
Up-regulation of mGluR1, ErbB4, and ErbB2 in SNpc DA neurons of the FXS mouse. **a)** Experimental pipeline for western blots and immunofluorescence experiments. **b)** Left, cropped representative western blots showing mGluR1 protein levels in total SNpc from WT (n = 7) and *Fmr1* KO mice (n = 10). The molecular weight of each protein is indicated in kDa. Right, quantification of the western blots presented in the left panel. The graphs show the quantification of the signal intensity normalized over the WT condition (%). (U = 11, P = 0.018, Mann-Whitney test). **c)** Cartoon depicting ErbB-dependent regulation of mGluR1 synthesis and membrane trafficking in SNpc DA neurons^33^. **d)** Left, cropped representative western blots showing ErbB4 protein levels in total SNpc from WT (n = 9) and *Fmr1* KO mice (n = 10). Right, quantification of the western blots presented in the left panel. The graphs show the quantification of the signal intensity normalized over the WT condition (%). (U = 19, P = 0.035, Mann-Whitney test). **e)** Left, cropped representative western blots showing ErbB2 protein levels in total SNpc from WT (n = 10) and *Fmr1* KO mice (n = 10). Right, quantification of the western blots presented in the left panel. The graphs show the quantification of the signal intensity normalized over the WT condition (%). (U = 27, P = 0.089. Mann-Whitney test). **f)** Left, cropped representative western blots showing NRG1 full-length and soluble forms protein levels in total SNpc from WT (n = 9) and *Fmr1* KO mice (n = 10). Right, quantification of the western blots presented in the left panel. The graphs show the quantification of the signal intensity normalized over the WT condition (%). (NRG1 full-length: U = 44.50, P = 0.983; NRG1 soluble form: U = 43, P = 0.887. Mann-Whitney test). **b,d-f)** Data are presented as mean ± SEM. Protein levels were normalized to the average of Coomassie staining, vinculin and β-actin. **g)** Left, double-labelled confocal images of mGluR1 (red) and tyrosine hydroxylase (TH) (green) in SNpc DA neurons captured at lower (up) and higher (bottom) magnification. Right, plot of mGluR1 integrated density (as % of WT) in nigral DA neurons of WT (n = 197 cells/3 mice) and *Fmr1* KO mice (n = 195 cells/3 mice). P = 0.037, U = 16871.5, Mann-Whitney test. **h)** Left, double-labelled confocal images of ErbB4 (red) and TH (green). Right, quantification of ErbB4 integrated density (normalized to WT) in SNpc DA neurons of WT (n = 257 cells/3 mice) and *Fmr1* KO (n = 392 cells/3 mice). P = 8.71 × 10^-20^, U = 29528, Mann-Whitney test. **i)** Left, double-labelled confocal images of ErbB2 (red) and TH (green) in SNpc DA neurons. Right, plot of ErbB2 integrated density (as % of WT) in nigral DA neurons from *Fmr1* KO (n = 200 cells/3 mice) and WT mice (n = 179 cells/3 mice). P = 1.859 × 10^-7^, U = 12403, Mann-Whitney test. **g-i)** Data are presented as mean ± SEM. Scale bar is 20 µm at lower magnification (up) and 10 µm at higher magnification (down).

What molecular event promotes mGluR1 up-regulation in the FXS mouse? We previously reported that the expression of mGluR1 in nigral DA neurons of wild-type rodents is regulated by ErbB tyrosine kinases, activated by the neurotrophic and differentiation factors neuregulins (NRGs)^33^. The NRGs family - including different types (NRG1-NRG6) and several splicing-derived isoforms - acts on homo- or heterodimeric receptors, composed of ErbB2, ErbB3, and ErbB4 subtypes^28^. ErbB4-ErbB4 and ErbB2-ErbB4 mediate NRGs/ErbB-induced effects on mGluR1 expression and function in SNpc DA neurons^33,34^. Dysregulation of the NRGs/ErbB axis has been firstly associated with schizophrenia, based on studies identifying *Nrg1* and *ErbB4* as risk genes and evidence linking NRGs/ErbB alterations with schizophrenic symptoms^35,36,37,38^. Recently, *Nrg1* polymorphisms have also been reported in ASD^39,40^. However, clinical evidence on NRG1 expression in the blood of patients with idiopathic ASD reported controversial data (describing increased^41,42^ or reduced levels^43^), thus the factual role of NRGs/ErbB signaling in ASD remains still elusive. Remarkably, NRGs/ErbB dysregulation has never been proposed in FXS.

Based on our previous evidence that ErbB stimulation promotes mGluR1 synthesis (**Fig. 3c**) and considering that *ErbB4* and *ErbB2* are listed as putative FMRP targets^44,45,46,47^, we predicted that FMRP deficiency triggers ErbB4 and ErbB2 up-regulation in nigral DA neurons and, subsequently, abnormal ErbB signaling promotes mGluR1 overproduction and neuronal hyperactivity. To test this hypothesis, we first checked the expression of nigral ErbB4 and ErbB2, revealing a robust increase in ErbB4 levels (**Fig. 3d**) and a trend toward up-regulation of ErbB2 in the FXS model (**Fig. 3e**). Differently, NRG1 – the prototypical ErbB ligand – either in a full-length membrane-anchored form or as a cleaved soluble domain appeared normally expressed in the FXS mouse (**Fig. 3f**). Next, to demonstrate that upregulations of mGluR1 and ErbB4/ErbB2 occur within nigral DA neurons, we quantified by immunofluorescence their optical densities in SNpc tyrosine hydroxylase positive (TH+) neurons. The results show increased expression of mGluR1 (**Fig. 3g**), ErbB4 (**Fig. 3h**), and ErbB2 (**Fig. 3i**) within nigral DA neurons in the FXS mouse.

### ErbB inhibition rescues nigral DA neuron dysfunctions in the FXS mouse

Next, we aimed to demonstrate that abnormal ErbB signaling drives mGluR1-dependent hyperactivity of SNpc DA neurons in the FXS mouse. Previously, we reported that tonic ErbB signaling shapes mGluR1 function in nigral DA neurons, by fine-tuning its membrane expression. The ErbB4 and ErbB2 inhibitor PD158780 hampers mGluR1 function by promoting receptor internalization^28,33,34^ (**Fig. 4a**). Here, we tested whether ErbB inhibition can attenuate mGluR1 hyperfunction and rescue hyperactivity and hyperexcitability of nigral DA neurons in the FXS mouse. We found that PD158780 (10 µM) reduced I_DHPG_ (**Fig. 4b, c**) and normalized spontaneous firing rate (**Fig. 4d**) and hyperexcitability (**Fig. 4e**) of nigral DA neurons of *Fmr1* KO mice. Collectively, these data support a causal link between amplified ErbB signaling, mGluR1 hyperfunction, and SNpc DA neuron hyperactivity. Furthermore, the identification of ErbB4/ErbB2 and mGluR1 dysfunction in the FXS model provides insights into the molecular mechanisms that go awry in FXS.

**Figure 4.**
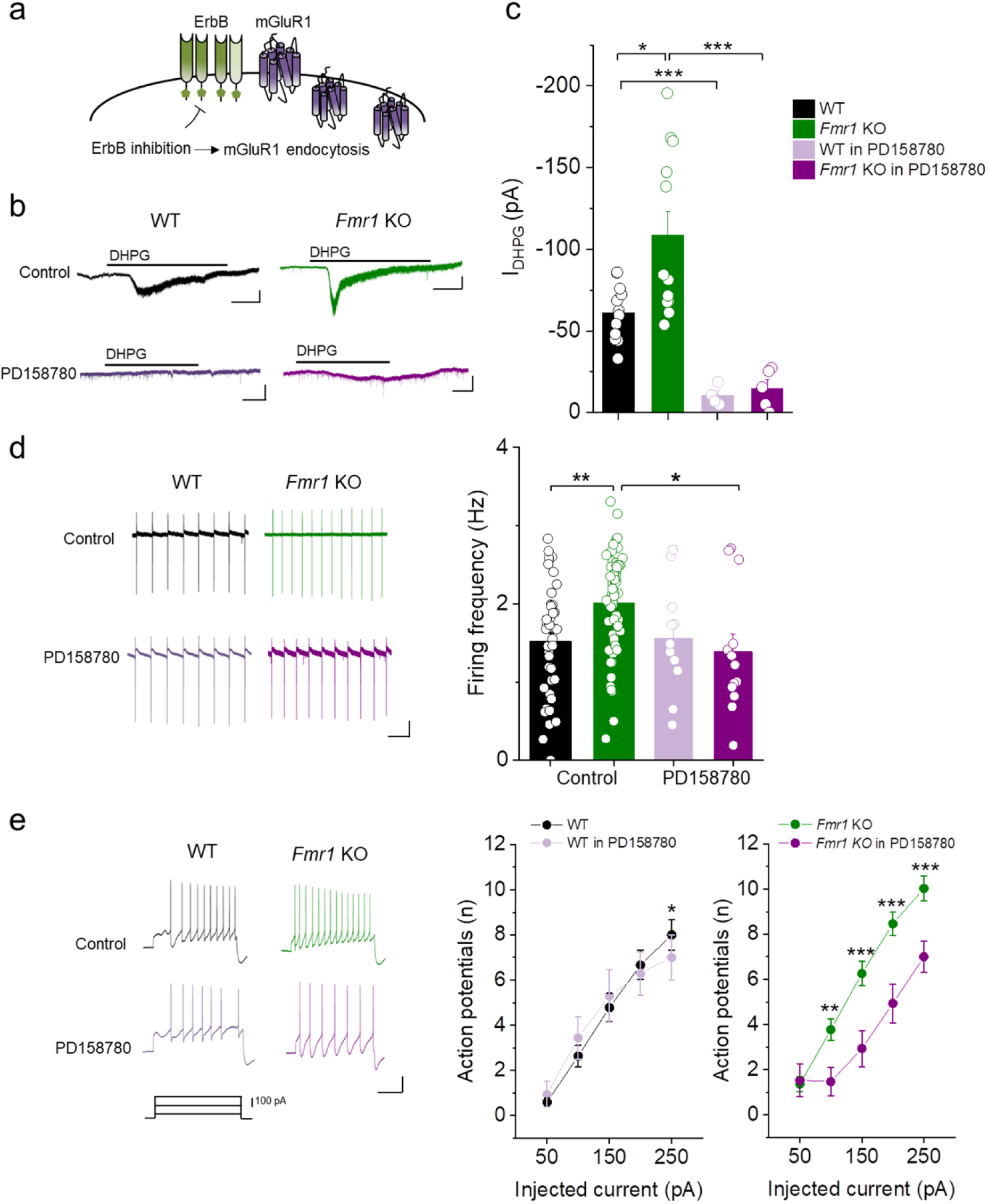
ErbB inhibition reduces mGluR1-dependent hyperactivity of SNpc DA neurons in the FXS mouse. **a)** Cartoon of ErbB-dependent regulation of mGluR1 expression in membranes showing that ErbB inhibition induces mGluR1 endocytosis in SNpc DA neurons^33^. **b)** Example I_DHPG_ traces in SNpc DA neurons in control and during treatment with PD158780 (10 µM). Scale bar: 20 pA/0.5 s. **c)** Plot of I_DHPG_ amplitudes showing that ErbB inhibition reduces mGluR1 function in both genotypes. WT-control (n = 12 cells/3 mice), *Fmr1* KO-control (n = 12 cells/4 mice), WT-PD158780 (n = 4 cells/3 mice), and *Fmr1* KO-PD158780 (n = 5 cells/3 mice). Kruskall-Wallis ANOVA followed by Mann-Whitney test. ANOVA P = 6.760 × 10^-5^. P = 0.020 for WT-control vs *Fmr1* KO-control; P = 0.0011 for WT-control vs WT-PD158780; P = 3.232 × 10^-4^ for *Fmr1* KO-control vs *Fmr1* KO-PD158780. **d)** Example firing traces (left) and quantification of firing frequency (right) in control and during PD158780 treatment. WT-control (n = 41 cells/8 mice), *Fmr1* KO-control (n = 59 cells/10 mice), WT-PD158780 (n = 11 cells/6 mice), and *Fmr1* KO-PD158780 (n = 13 cells/6 mice). Two-way ANOVA followed by Tukey’s test. Genotype x drug interaction P=0.0348; P=0.002 for WT vs *Fmr1* KO, P=0.015 *Fmr1* KO vs *Fmr1* KO in PD158780. Scale bar: 20 pA/1 s. **e)** Example traces (left) and plots (right) of depolarizing-evoked APs, showing that ErbB inhibition normalizes SNpc DA neuron hyperexcitability in the FXS mouse. PD158780 effect in WT: WT-control (n = 27 cells/7 mice) and WT-PD15870 (n = 14 cells/5 mice). RM two-way ANOVA followed by Tukey’s test. Drug P=0.618, injected current P = 7.413 X10^-28^, drug X injected current P = 0.032. P = 0.0341 at 250 pA. PD158780 effect in *Fmr1* KO: *Fmr1* KO-control (n = 39 cells/10 mice) and *Fmr1* KO-PD158780 (n = 15 cells/6 mice). Drug P = 0.006, injected current P = 1.271 × 10^-28^, drug x injected current P = 1.514 × 10^-4^. P = 0.003 at 100 pA, P = 1.735 × 10^-5^ at 150 pA, P = 1.089 × 10^-5^ at 200 pA, P = 6.825 × 10^-5^ at 250 pA. Scale bar: 25 mV/0.5 s. All data are presented as mean ± SEM.

### Inhibition of nigral ErbB *in vivo* normalizes SNpc DA neuron activity and repetitive behaviors in the FXS mouse

ASD-like behavioral alterations, including repetitive compulsive behaviors, have been consistently reported in adult *Fmr1* KO mice, with minor variances based on genetic background^48,49,50,51,52,53,54^. Differently, adolescent *Fmr1* KO mice have been rarely examined, therefore there is still an inadequate understanding of early behavioral alterations, which better reflect pediatric symptoms of FXS. To advance this knowledge, we evaluated whether adolescent 30-day-old *Fmr1* KO mice displayed abnormal repetitive behaviors (**Fig. 5a,b**). We observed a selective exacerbation of self-grooming (**Fig. 5c**), and normal digging (**Fig. 5d**) and jumping behavior (**Fig. 5e**) in the FXS mouse. Furthermore, in the marble burying test (also estimating repetitive behaviors) adolescent *Fmr1* KO mice reached higher scores than WT (**Fig. 5f**). These findings demonstrate early appearance of abnormal repetitive behaviors in the FXS model.

**Figure 5.**
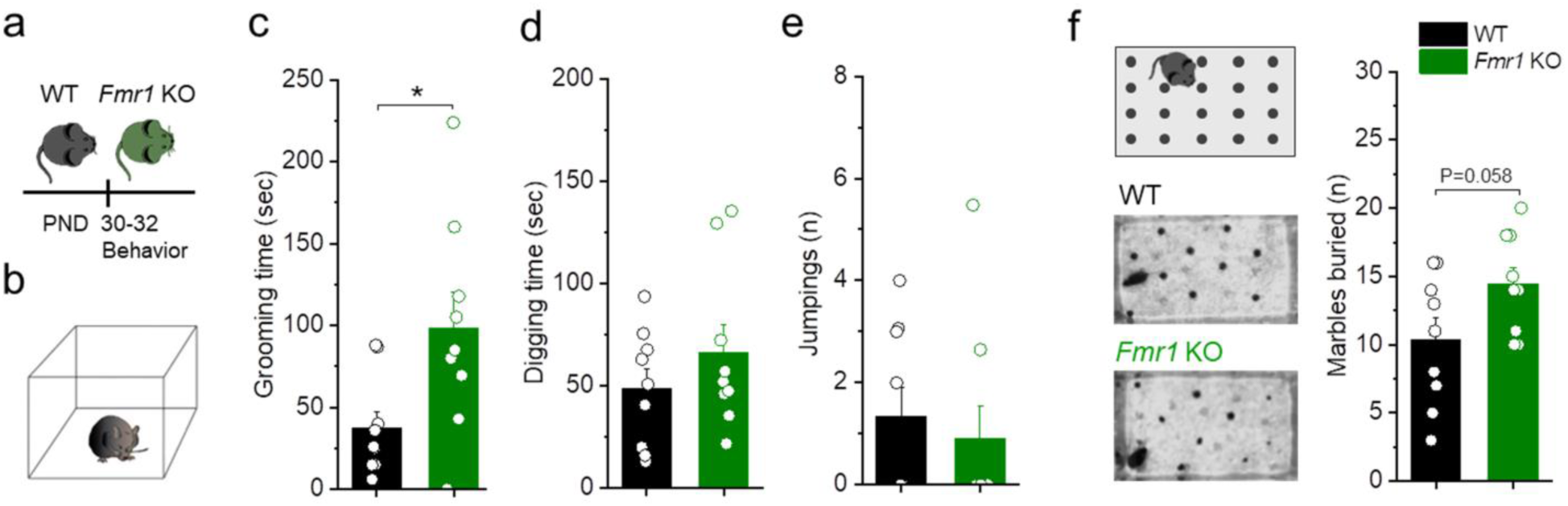
Early appearance of abnormal repetitive behaviors in adolescent 30-day-old *Fmr1* KO mice. **a)** Timeline of behavioral experiments; **b)** Setting for analyses of spontaneous repetitive behaviors and quantifications of time spent in self-grooming **(c)** or digging **(d)** and number of jumping **(e)** by *Fmr1* KO (n = 9) and WT (n = 9) mice. Adolescent *Fmr1* KO mice exhibit a selective exacerbation of self-grooming. P = 0.038, U = 17, Mann-Whitney test. Digging time: P = 0.302, two-tailed unpaired *t*-test. Jumping (n): P = 0.612, Mann-Whitney test. **f)** Left, Schematic of marble burying test and example images of tasks executed by WT and *Fmr1* KO mice; Right, Quantification of marbles buried showing increased compulsive burying in the FXS mouse (n = 9 mice for each group, P = 0.058, *t* = -2.040, two-tailed unpaired *t*-test. All data are presented as mean ± SEM.

Then, we predicted that abnormal ErbB signaling, by shaping nigral DA neurons activity, would drive compulsive repetitive behaviors in the FXS model. Consistent with this, *in vivo* inhibition of nigral ErbB signaling should simultaneously rescue nigral DA neuron hyperactivity and repetitive behaviors in the FXS mouse. To assess this hypothesis we first evaluated the effects of *in vivo* nigral ErbB inhibition on SNpc DA neuron dysfunctions. *Fmr1* KO and WT mice were subjected to cannulation to perform bilateral intra-SNpc injections of PD158780 (10 µM, 0.6 µL) or its vehicle (Veh,0.1 % DMSO), followed by patch clamp recordings of nigral DA neurons in midbrain slices of the injected mice (**Fig. 6a**). We found that nigral ErbB inhibition lowers I_DHPG_ (**Fig. 6b**) and corrects spontaneous firing hyperactivity (**Fig. 6c**) and hyperexcitability (**Fig. 6d**) of SNpc DA neurons of the FXS mouse. Thus, *in vivo* intra-SNpc ErbB inhibition, by offsetting mGluR1 hyperfunction, normalizes the activity of nigral DA neurons in the FXS model.

**Figure 6.**
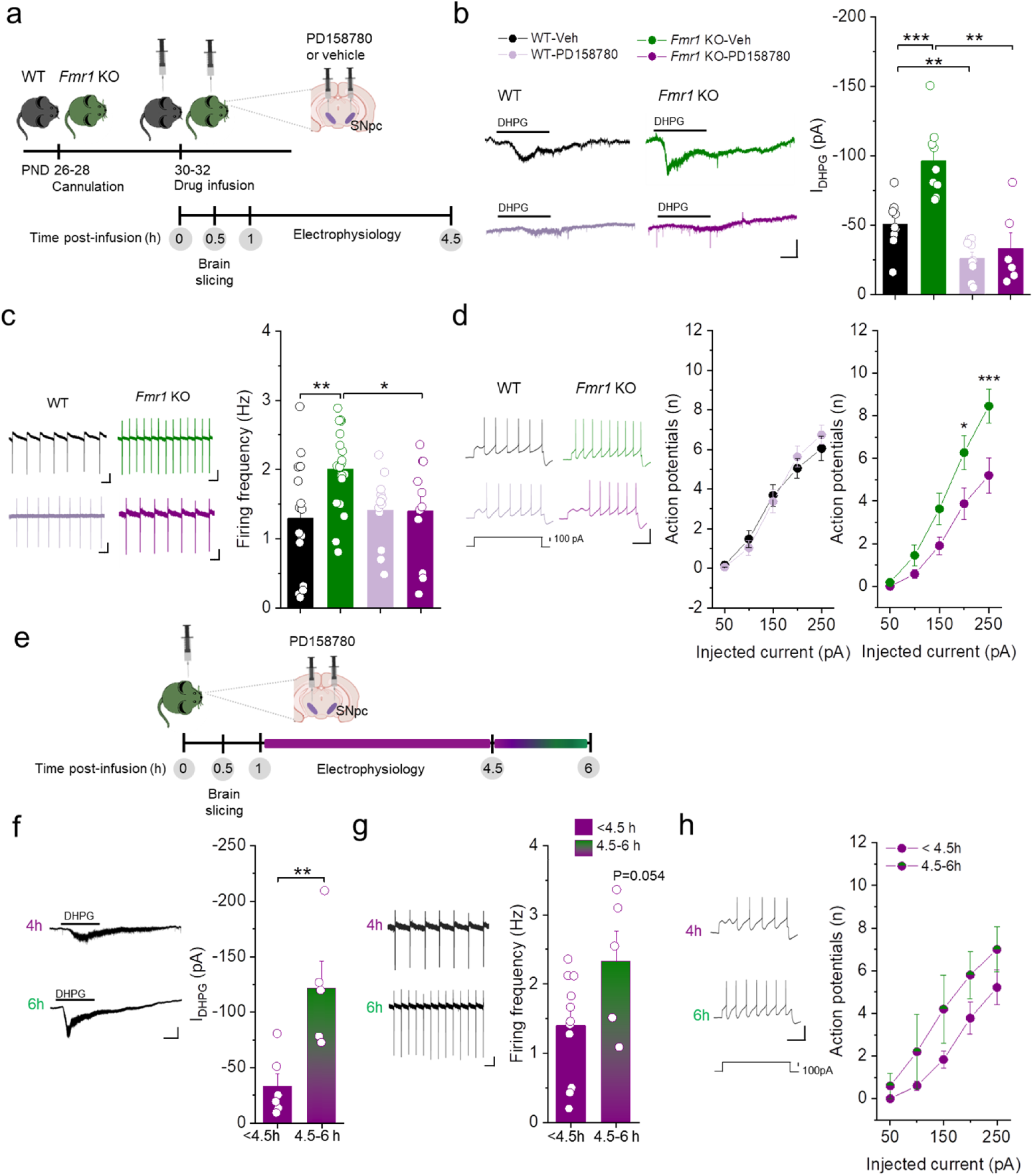
Inhibition of nigral ErbB *in vivo* normalizes SNpc DA neuron functions in the FXS mouse. **a)** Experimental design to evaluate the effects of nigral ErbB inhibition on SNpc DA neuron dysfunctions in the FXS model. Adolescent mice were subjected to intra-SNpc injections of PD158780 (10 µM) or its vehicle (0.1 % DMSO in aCSF) before *ex vivo* electrophysiology. **b)** Example I_DHPG_ traces (left) and plot of I_DHPG_ amplitude (right) showing local ErbB inhibition reduces I_DHPG_ in both genotypes. WT-Vehicle (n = 9 cells/3 mice), WT-PD158780 (n = 9 cells/3 mice), *Fmr1* KO-Vehicle (n = 9 cells/3 mice), and *Fmr1* KO-PD158780 (n = 6 cells/3 mice). Kruskall-Wallis ANOVA followed by Mann-Whitney test. P = 1.234 × 10^-4^ for ANOVA; P = 2.879 × 10^-4^ for WT-Vehicle vs *Fmr1* KO-Vehicle, P = 0.003 for WT-Vehicle vs WT-PD158780, P = 0.003 for *Fmr1* KO-Vehicle vs *Fmr1* KO-PD158780. Scale bar: 50 pA/1 min. **c)** Example firing traces (left) and plot of firing frequency (right) proving that nigral ErbB inhibition decreases SNpc DA neuron hyperactivity of the FXS mouse. WT-Vehicle (n = 15 cells/3 mice), WT-PD158780 (n = 12 cells/3 mice), *Fmr1* KO-Vehicle (n = 20 cells/3 mice), and *Fmr1* KO-PD158780 (n = 11 cells/3 mice). Two-way ANOVA followed by Fisher’s test. ANOVA: genotype P=0.059, drug P = 0.179, genotype X drug interaction P = 0.048. P = 0.002 for WT-Vehicle vs *Fmr1* KO-Vehicle, P = 0.018 for WT-PD158780 vs *Fmr1* KO-Vehicle, P = 0.018 for *Fmr1* KO-Vehicle vs *Fmr1* KO-PD158780. Scale bar: 20pA/0.5 s. **d)** Representative evoked APs traces (left) and plot of APs number (right) showing ErbB inhibition offsets hyperexcitability of SNpc DA neurons in the FXS model. WT-Vehicle (n = 19 cells/3 mice) and WT-PD158780 (n = 19 cells/3 mice), RM two-way ANOVA. Interaction P = 0.440. *Fmr1* KO-Vehicle (n = 22 cells/3 mice) and *Fmr1* KO-PD158780 (n = 18 cells/3 mice), RM two-way ANOVA followed by Tukey’s test. ANOVA: drug P = 0.073, drug X current interaction P = 0.015. P = 0.018 at 200 pA, P = 5.41 X10^-4^ at 250 pA. Scale bar: 20 mV/0.5 s. **e)** Timeline of experiments to analyze the kinetics of ErbB inhibition-induced effects in the FXS mouse. **f)** Example I_DHPG_ traces (left) and plot of I_DHPG_ amplitude (right) showing initial rescues of mGluR1 function at 4.5-6 h after intra-SNpc PD158780 injection. I_DHPG_ < 4.5 h (n = 6 cells/3 mice) and I_DHPG_ at 4.5-6 h (n = 5 cells/3 mice); P=0.007, t= 3.474, two-tailed unpaired *t-*test. Scale bar: 50 pA/1 min. **g)** Representative firing traces (left) and plot of firing frequency (right) of SNpc DA neurons recorded at < 4.5 h (n = 11 cells/3 mice) or 4.5-6 h (n = 5 cells/3 mice) after nigral ErbB inhibition. A trend toward an increase in spontaneous firing frequency is overt at 4.5-6h after local PD158780 injection. P = 0.054, t =-2.100, two-tailed unpaired *t*-test. Scale bar: 20 pA/0.5 s. **h)** Evoked APs traces (left) and APs number (right) showing a trend, still not significant, toward neuronal hyperexcitability in the FXS mouse at 4.5-6h after PD158780 injection. Post-injection time: < 4.5 h (n = 18 cells/3 mice) and 4.5-6 h (n = 5 cells/3 mice). RM two-way ANOVA. Time P = 0.202, time X current interaction P = 0.865. Scale bar: 20 pA/0.5 s. All data are presented as mean ± SEM.

To estimate the duration of rescue effects induced by nigral ErbB inhibition, we examined the reemergence of SNpc DA neuron dysfunctions in the FXS mouse after intra-SNpc injection of PD158780 (**Fig. 6e**). SNpc DA neuron dysfunctions are completely neutralized for at least 4.5 h after intra-SNpc injection of the ErbB inhibitor; after that, at 4.5-6 h after injection, there is partial regain of I_DHPG_ (**Fig. 6f**) and a trend toward nigral DA neuron hyperactivity (**Fig. 6g**) and hyperexcitability (**Fig. 6h**).

To evaluate the effects of the inhibition of nigral ErbB on repetitive behaviors, we examined self-grooming and marble burying behavior in mice subjected to bilateral infusion of the ErbB inhibitor or its vehicle in the SNpc (**Fig. 7a**). We found that nigral ErbB inhibition rescues exacerbated self-grooming (**Fig. 7b**) and improves marbles burying behavior of *Fmr1* KO mice (**Fig. 7c**). General locomotor activity of *Fmr1* KO mice in an open field was not affected **(Supplementary Fig. 4).** This evidence indicates specific reduction of repetitive behaviors of the FXS mouse. Collectively, our findings demonstrate that ErbB inhibition in the SNpc concomitantly corrects dysfunctions of nigral DA neurons and aberrant repetitive behaviors in the FXS model.

**Figure 7.**
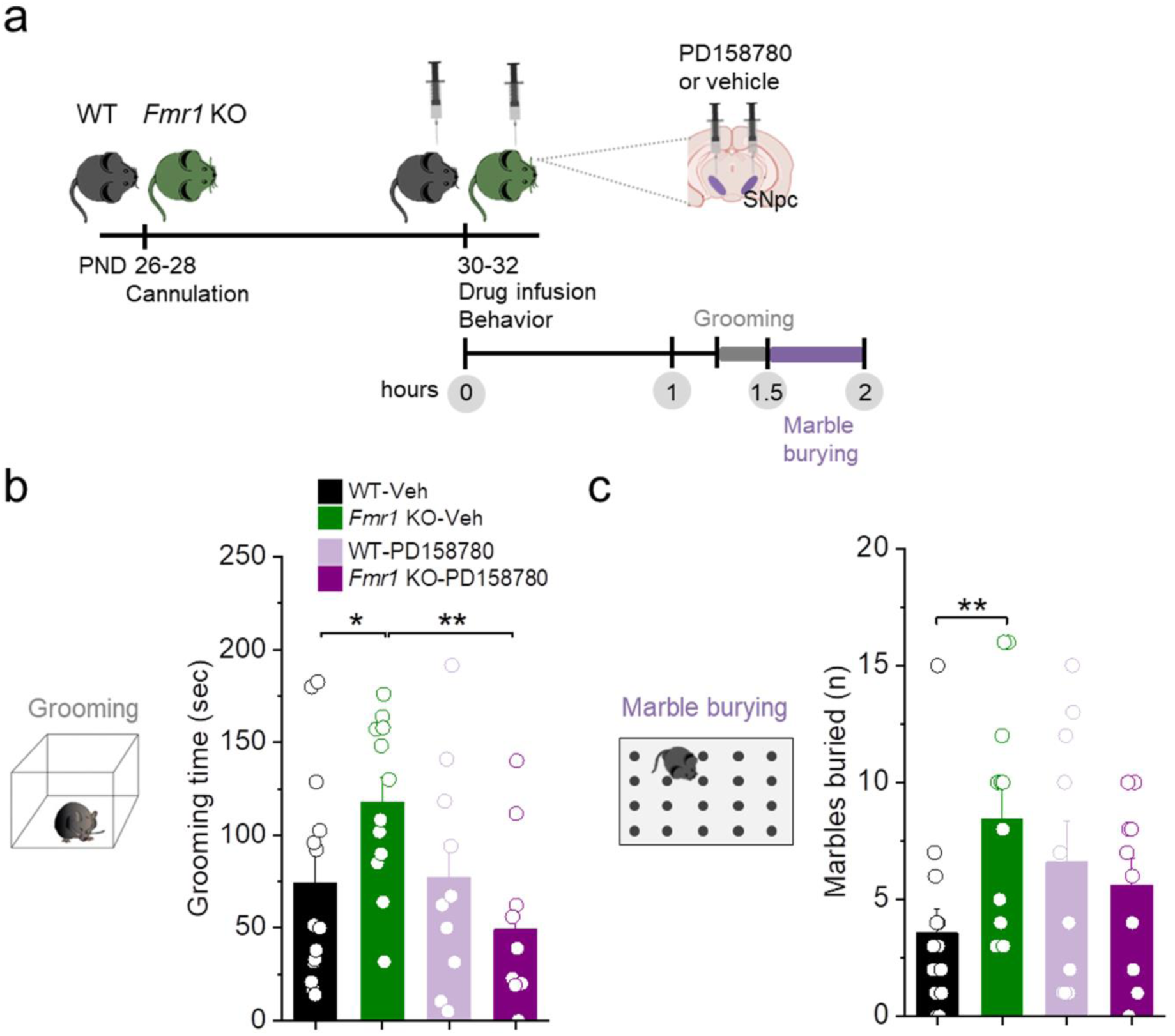
Nigral ErbB inhibition reduces abnormal repetitive behaviors in the FXS mouse. **a)** Experimental design to evaluate the effects of ErbB inhibition in the SNpc on repetitive behaviors of adolescent mice. **b)** Plot of time spent in self-grooming showing that nigral ErbB inhibition rescues this repetitive behavior in the FXS mouse. WT-Vehicle (n = 14), WT-PD158780 (n = 10), *Fmr1* KO-Vehicle (n = 12), and *Fmr1* KO-PD158780 (n = 10). Two-way ANOVA followed by Fisher’s test. ANOVA: drug P = 0.041, genotype X drug interaction P = 0.026. P = 0.040 for WT-Vehicle *vs Fmr1* KO-Vehicle; P = 0.003 for *Fmr1* KO-Vehicle vs *Fmr1* KO-PD158780. **c)** Number of marbles buried indicating that nigral ErbB inhibition attenuates marble burying in the FXS model. WT-Vehicle (n = 14), WT-PD158780 (n=10), *Fmr1* KO-Vehicle (n = 12), and *Fmr1* KO-PD158780 (n = 10). Two-way ANOVA followed by Fisher’s test. ANOVA: genotype X drug interaction P = 0.033. P = 0.008 for WT-vehicle vs *Fmr1* KO-vehicle, P=0.147 for *Fmr1* KO-vehicle vs *Fmr1* KO-PD158780.

### Systemic ErbB inhibition corrects SNpc DA neuron dysfunctions and repetitive behaviors of the FXS mouse

To understand if ErbB targeting is a feasible approach for FXS treatment, we examined whether the systemic administration of the ErbB inhibitor is equally effective as intra-SNpc ErbB inhibition in rescuing SNpc DA neuron hyperactivity and repetitive behaviors of the FXS model. Adolescent *Fmr1* KO and WT mice were injected intraperitoneally with PD158780 (10 mg/Kg) or its vehicle (30% DMSO in 0.9 % NaCl) and then were subjected to procedures for electrophysiological or behavioral studies (**Fig. 8a**). First, we performed patch clamp recordings of SNpc DA neurons in midbrain slices from vehicle- or PD158780-treated mice and found that systemic ErbB inhibition mimics the effects of nigral ErbB inhibition, rescuing SNpc DA neuron dysfunctions in the FXS model. Precisely, PD158780-treated *Fmr1* KO mice showed reduced I_DHPG_ (**Fig. 8b**) and normal spontaneous firing frequency (**Fig. 8c**) and excitability (**Fig. 8d**), demonstrating the efficacy of systemic ErbB inhibition in compensating for nigral DA neuron hyperactivity.

**Figure 8.**
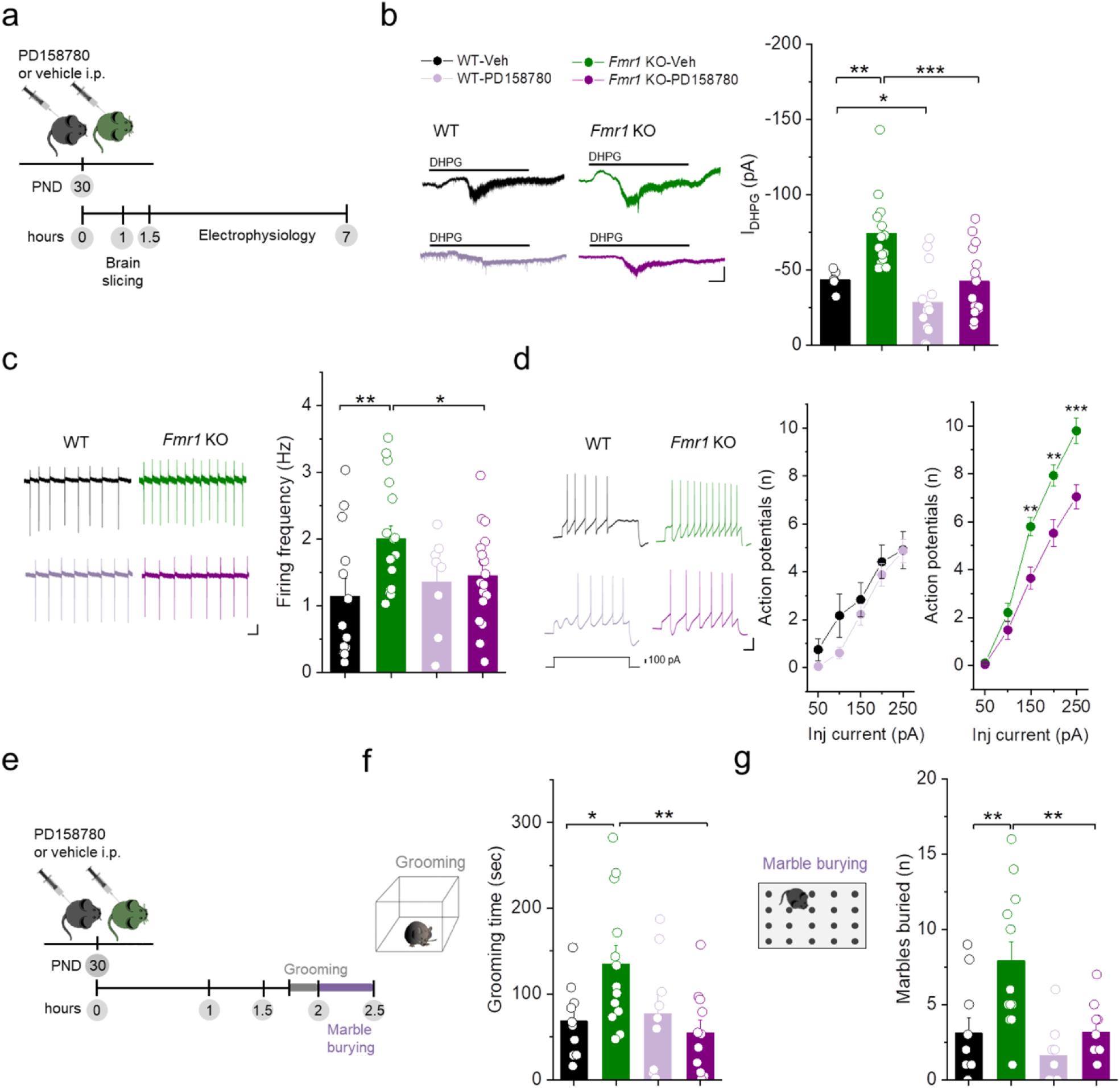
Systemic ErbB inhibition corrects SNpc DA neuron dysfunctions and repetitive behaviors of the FXS mouse. **a)** Experimental design to evaluate the effects of systemic ErbB inhibition on SNpc DA neuron functions. **b)** Example I_DHPG_ traces (left) and quantification of I_DHPG_ amplitude (right) in WT-Vehicle (n = 5 cells/2 mice), WT-PD158780 (n = 13 cells/3 mice), *Fmr1* KO-Vehicle (n = 14 cells/3 mice), and *Fmr1* KO-PD158780 (n = 15 cells/3 mice). Kruskal-Wallis ANOVA followed by Kolmogorov-Smirnov test. ANOVA: P = 0.001. P = 0.039 for WT-Vehicle vs WT-PD158780, P = 0.001 for WT-Vehicle vs *Fmr1* KO-Vehicle, P = 8.123 × 10^-4^ for *Fmr1* KO-Vehicle vs *Fmr1* KO-PD158780. Scale bar: 50 pA/0.5 s. **c)** Example spontaneous firing traces (left) and firing frequency plot (right) of WT-Vehicle (n = 13 cells/2 mice), WT-PD158780 (n = 8 cells/3 mice), *Fmr1* KO-Vehicle (n = 17 cells/3 mice), and *Fmr1* KO-PD158780 (n = 18 cells/3 mice). Two-way ANOVA followed by Fisher’s test. ANOVA: genotype P = 0.038, genotype X drug interaction P = 0.093. P = 0.005 for WT-Vehicle vs *Fmr1* KO-Vehicle, P = 0.045 *Fmr1* KO-Vehicle vs *Fmr1* KO-PD158780. Scale bar: 10 pA/0.5 s. **d)** Evoked APs traces (left) and APs number plot (right). Effects in WT mice: WT-Vehicle (n = 12 cells/2 mice) and WT-PD158780 (n = 23 cells/3 mice). RM two-way ANOVA. Drug P = 0.393, drug X current interaction P = 0.768. Effect in *Fmr1* KO mice: *Fmr1* KO-Vehicle (n = 29 cells/3 mice) and *Fmr1* KO-PD158780 (n = 25 cells/3 mice). RM two-way ANOVA followed by Bonferroni test. ANOVA: drug P = 1.76 × 10^-4^, drug X current interaction P = 2.699 × 10^-4^. P = 0.009 at 150 pA, P = 0.003 at 200 pA, P = 3.003 × 10^-4^. Scale bar: 20 mV/0.5. **e)** Experimental protocol to analyze the effects of systemic ErbB inhibition on aberrant repetitive behaviors in adolescent FXS mice. **f)** Plot of time spent in self-grooming by WT-Vehicle (n = 10), WT-PD158780 (n = 9), *Fmr1* KO-Vehicle (n = 13), and *Fmr1* KO-PD158780 (n = 11). Two-way ANOVA followed by Fisher’s test. ANOVA: genotype X drug interaction P = 0.023. P = 0.012 for WT-Vehicle vs *Fmr1* KO-Vehicle, P = 0.002 for *Fmr1* KO-PD158780 vs *Fmr1* KO-Vehicle. **g)** Number of marbles buried by WT-Vehicle (n = 10), WT-PD158780 (n = 9), *Fmr1* KO-Vehicle (n = 13) and *Fmr1* KO-PD158780 (n = 11). Two-way ANOVA followed by Bonferroni test. ANOVA: genotype P = 0.001, drug P = 0.002, genotype X drug interaction P = 0.099. P = 0.004 for WT-Vehicle vs *Fmr1* KO-Vehicle, P = 0.004 for F*mr1* KO-Vehicle vs *Fmr1* KO-PD158780.

In parallel, we analyzed the effects of systemic inhibition of ErbB signaling on abnormal repetitive behaviors of the FXS mouse (**Fig. 8e**). We found that systemic administration of the ErbB inhibitor rectifies compulsive self-grooming (**Fig. 8f**) and marble burying behavior of adolescent *Fmr1* KO mice (**Fig. 8g**), without affecting their general locomotor activity **(Supplementary Fig. 5).** Thus, systemic ErbB inhibition recapitulates rescue effects reliant on inhibition of ErbB in the SNpc. Altogether, these findings demonstrate that systemic administration of the ErbB inhibitor restores normal SNpc DA neuron activity and regular behaviors in the FXS model.

In conclusion, the present manuscript outlines a previously unrealized mechanism contributing to FXS (**Fig. 9**). In the FXS model, loss of FMRP in nigral DA neurons drives up-regulation of ErbB4 and ErbB2, subsequently promoting mGluR1 overproduction; in turn, exacerbated ErbB/mGluR1 function enhances firing activity of SNpc DA neurons, fostering the emergence of repetitive behaviors. Consistently, ErbB inhibition in the FXS mouse, by mitigating abnormal mGluR1 tone, neutralizes nigral DA neuron hyperactivity and repetitive behaviors. Collectively, our evidence demonstrates that SNpc DA neuron hyperactivity is an early signature of FXS, nigral ErbB4/ErbB2 and mGluR1 play a relevant role in the etiology of FXS, and targeting ErbB represents a valuable pharmacological approach to treat the core symptoms of ASD and FXS.

**Figure 9.**
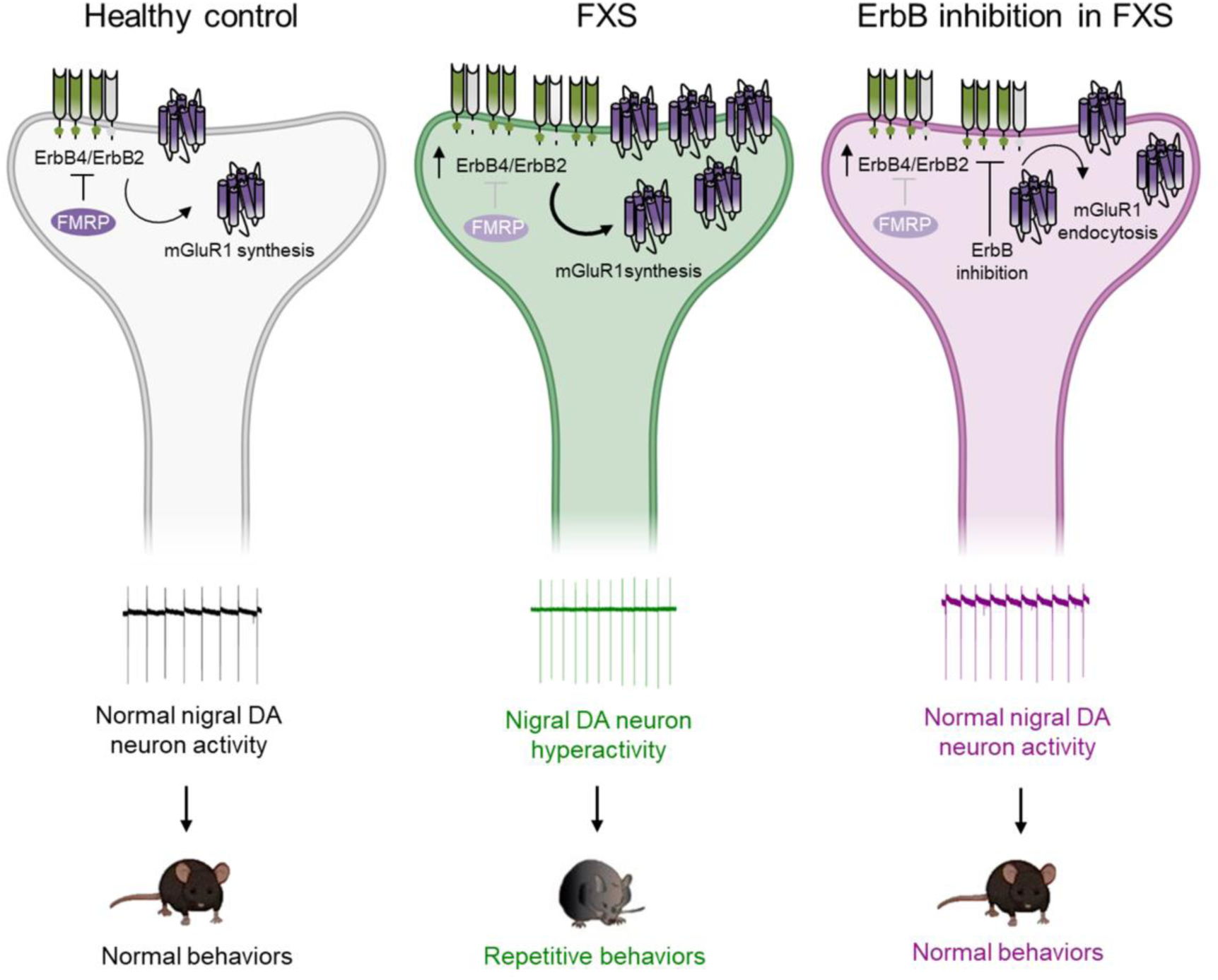
Hyperfunction of ErbB4/ErbB2 and mGluR1 drives SNpc DA neuron hyperactivity and repetitive behaviors in the FXS model. Left) In the healthy brain, ErbB4/ErbB2 signaling fine-tunes mGluR1 expression and function in nigral DA neurons, contributing to keep firing activity at physiological levels instrumental to normal behaviors. Middle) In the FXS model, FMRP deficiency induces ErbB4 and ErbB2 up-regulation and consequential mGluR1 overproduction in nigral DA neurons. This exacerbated ErbB/mGluR1 function induces neuronal hyperactivity and hyperexcitability, leading to abnormal activation of nigrostriatal DA circuit and insurgence of repetitive behaviors. Right) ErbB inhibition in the FXS model, by triggering mGluR1 endocytosis, neutralizes aberrant ErbB/mGluR1-dependent effects within nigral DA neurons, restoring normal firing activity and regular behaviors.

## Discussion

Beyond cognitive disability, FXS is characterized by a complex symptomatology, including epilepsy, auditory hypersensitivity, locomotor hyperactivity, anxiety, social avoidance, and repetitive stereotyped behaviors, with a high incidence of autism, diagnosed in approximately 50% of males and 20% of females with FXS^2,55,56^. Multifaceted symptomatology advises the contribution of multiple brain areas; indeed, functional alterations have been described in the hippocampus, cortex, thalamus, cerebellum, amygdala, and striatum of FXS models and patients^3,27,56^. Although the SNpc DA nucleus is a central hub for movement control and the creation of habits and sequential behaviors, and it is expected that its dysfunction could trigger compulsive and stereotyped behaviors^14,15^, research aimed at identifying dysfunctions of nigral DA neurons in ASD and FSX models has been neglected so far. Here, we provide the first direct evidence of early dysregulation of the SNpc DA nucleus in FXS, revealing various functional changes that occur in nigral DA neurons of the FXS mouse in adolescence since the third postnatal weeks. Precisely, SNpc DA neurons in the FXS mouse display an increased spontaneous firing rate and hyperexcitability, in addition to alterations in passive membrane properties, namely increased C_m_. Since C_m_ depends on membrane size and composition, higher C_m_ in the FXS mouse may reflect hypertrophic changes in the soma or dendritic arbor and/or altered lipidic double-bilayer composition or thickness. Notably, increased density of TH^+^ fibers in the striatum of 3-month-old *Fmr1* KO has been recently described, suggesting hypertrophic changes in nigral DA neurons in adulthood^20^.

Neuronal hyperexcitability is a recurrent signature in FXS, associated with abnormal circuit connectivity in different brain areas, including the somatosensory and entorhinal cortex, auditory brainstem, and hippocampus^27^. Diverse and apparently area-specific underlying mechanisms have been identified, including E/I imbalance (i.e., increased glutamatergic or reduced GABAergic transmission), or dysfunctions in different types of K^+^ and Na^+^ channels, such as voltage-dependent Kv3.1 or Ca^2+^-activated K^+^ channels (SK and BK) or Na^+^ channels (Slack, and those carrying persistent or transient Na^+^ currents, I_NaP_ and I_NaT_)^27,57,58,59,60,61,62,63^. Ion channels alterations can directly arise from increased translation or loss of physical FMRP-channels interaction; otherwise, they can be secondary to abnormal expression/activity of upstream regulators, as reported for mGluR5^60,64^. The present manuscript advances the understanding of FXS substrates by revealing that mGluR1 hyperfunction drives nigral DA neuron hyperactivity and hyperexcitability in the FXS model. In addition to directly evidencing higher mGluR1-induced currents (I_DHPG_ in MPEP), we show that mGluR1 inhibition normalizes the spontaneous firing rate and excitability of SNpc DA neurons in the FXS mouse.

Knowledge about mGluR1 role in FXS etiology is rather limited: some evidence indicates that mGluR1 inhibition, by antagonists or genetic reduction, ameliorates behavioral phenotypes such as audiogenic seizures, hyperlocomotion, and marble burying in adult *Fmr1* KO mice^65,66^, but precise information on brain areas, cellular loci, and specific mGluR1-dependent mechanism contributing to FXS is lacking. Contrariwise, mGluR5 has been in the spotlight of intense investigations in the FXS field since decades: exacerbated mGluR5 function is considered an FXS hallmark and motivated the “mGluR theory of FXS”^67^, which assumes that mGluR5 hyperfunction determines synaptic, behavioral, and cognitive FXS alterations. However, while mGluR5 inhibition attenuates some FXS phenotypes in preclinical models^68,60,70^, mGluR5-based therapeutics returned with limited efficacy in human clinical trials^71^, exposing the need to identify additional targets or collateral mechanisms beyond mGluR5 hyperfunction. Our data support the implication of mGluR1 in FXS etiology, proving its key role in influencing nigrostriatal DA circuit. Interestingly, a recent study attested association of *Grm1* (encoding for mGluR1) with schizophrenia^72^; therefore, mGluR1 dysfunction seems to be a shared feature of schizophrenia and FXS.

Different mechanisms could underlie mGluR1 hyperfunction in nigral DA neurons of the FXS model, that is, 1) increased glutamate release, acting on the same mGluR1 pool, 2) enhanced mGluR1 levels in the membrane, with or without variation in ambient glutamate, or 3) abnormal constitutive activity (ligand-independent signaling) of mGluR1 associated or not with changes in its levels. Analyses of glutamatergic synaptic inputs to nigral DA neurons exclude that increased glutamate release promotes mGluR1 hyperfunction and nigral DA neuron hyperactivity. In addition, normal GABAergic synaptic inputs similarly indicate that neuronal hyperactivity is not reliant on synaptic disinhibition. Diversely, we found higher mGluR1 levels in total protein extracts from SNpc homogenates and an increase in mGluR1 optical density in SNpc TH^+^ neurons of adolescent FXS mice, which demonstrate mGluR1 up-regulation within SNpc DA neurons. Therefore, an enrichment of the membrane-expressed mGluR1 pool, despite regular glutamate release, maintains mGluR1 hyperfunction and nigral DA neuron hyperactivity in the FXS mouse.

We have previously reported that ErbB signaling controls mGluR1 expression and function within SNpc DA neurons of WT rodents^28,33,34,73^. Precisely, NRG-activated ErbB signaling induces *de novo* synthesis and trafficking of mGluR1 to membranes, potentiating mGluR1-dependent effects, such as I_DHPG_, but also long-term depression^33,34^. Contrariwise, ErbB inhibition stimulates mGluR1 endocytosis in SNpc DA neurons, promptly disrupting all mGluR1 functions^28,33,34,73^. In light of this functional liaison between ErbB and mGluR1, and since *ErbB4* and *ErbB2* are putative FMRP targets^44,45,46,47^, we hypothesized that ErbB4/ErbB2 up-regulation may cause abnormal *de novo* synthesis of mGluR1 inside nigral DA neurons. Coherently, ErbB4 and ErbB2 levels, besides mGluR1, are increased in SNpc DA neurons, and with up-regulation gradients (ErbB4 and ErbB2 > mGluR1) supporting a hierarchic relationship. This supports an amplification of ErbB-induced mGluR1 synthesis^33^ in the FXS model.

NRGs/ErbB signaling is essential for correct neurodevelopment, by affecting neuronal migration, axon guidance, glia differentiation, myelination, neurite outgrowth, and synapses formation^74^, as well as for the regulation of adult brain functions, by shaping synaptic transmission and plasticity^73^. NRGs/ErbB dysregulation has recently been correlated with ASD, based on evidence of *NRG1* polymorphisms^39,40^ and altered NRG1 levels in blood from patients with idiopathic ASD^41,42,43^. Currently, preclinical analyses of NRGs/ErbB pathway in ASD models are restricted to a report showing increased levels of the NRGs subtypes (NRG3 and NRG1 type II) in microglia of BTBR mice (an idiopathic ASD model)^41^, and a second study analyzing ErbB inhibition-induced effects on hippocampal synaptic plasticity and contextual fear memory in a model of Angelman syndrome^75^. Due to this limited evidence, the factual contribution of the NRGs/ErbB system to ASD remains elusive. Remarkably, despite its recognized function in neurodevelopment, NRGs/ErbB signaling has not been previously evaluated in FXS. Therefore, the present demonstration of up-regulation of ErbB4 and ErbB2 in nigral DA neurons of the FXS mouse is the first evidence of NRGs/ErbB alteration in FXS. Notably, NRG1 expression in SNpc is regular in adolescent *Fmr1* KO mice. While ErbB4/ErbB2 upregulation *per se* can sustain enhanced ErbB signaling, despite normal amounts of ligands, we do not exclude that other NRGs subtypes beyond NRG1 could be up-regulated and future investigations might reveal these variations. Regardless of the precise ligand, our data prove that ErbB4/ErbB2 upregulation during critical developmental stages alters nigrostriatal DA circuit activity; counteracting nigral ErbB signaling, by *in situ* injection of an ErbB4 and ErbB2 inhibitor, besides blunting mGluR1-activated currents, normalizes spontaneous firing rate and excitability of SNpc DA neurons. Thus, tuning down ErbB signaling in the FXS mouse rescues nigral DA neuron functions.

We have previously reported that *in vivo* activation of nigral mGluR1 increases striatal DA release in rats, and this can be prevented by inhibiting ErbB^33^. Based on this evidence and previous indications that striatal DA tone affects repetitive behaviors in wild-type rodents, we predicted that ErbB/mGluR1-dependent nigral DA hyperactivity triggers repetitive behaviors in the FXS model. Soundly with this idea, exacerbated repetitive behaviors (self-grooming and marble burying) are overt in the FXS mouse at 30 days of age, a time-point at which nigral DA neurons hyperactivity is well-established. Importantly, this evidence describes the earliest appearance of ASD-like repetitive behaviors in *Fmr1* KO mice, previously reported in older animals (> 7 weeks)^50,52,54,76^.

According to our prediction, *in vivo* nigral ErbB inhibition concomitantly rescues functional alterations of SNpc DA neurons (by decreasing I_DHPG_ and normalizing spontaneous firing rate and excitability) and repetitive behaviors (by reducing self-grooming and marble burying). This proves a causal role for abnormal ErbB signaling, by activating the SNpc DA nucleus, in triggering repetitive behaviors in the FXS mouse. Consistent with this concept that ErbB overstimulation in early developmental stages promotes nigral DA neuron hyperactivity, a previous study in wild-type mice shows that subchronic neonatal exposure to NRG1 (from P2-P10) increases the spontaneous firing frequency of SNpc DA neurons in adulthood^77^.

Importantly, systemic administration of the ErbB inhibitor in the FXS mouse mimics the rescue effects induced by nigral ErbB inhibition, by rectifying all the functional anomalies that occur in SNpc DA neurons (higher mGluR1-induced currents, hyperactivity, and hyperexcitability), as well as abnormal self-grooming and marble burying. Thus, besides revealing a novel mechanism - ErbB/mGluR1-regulated SNpc DA neuron hyperactivity - instrumental to repetitive behaviors, the present study demonstrates that ErbB inhibition, also via a practicable route of administration, effectively reduces FXS symptoms.

Beyond unveiling the functional events associated with ErbB inhibition in the FXS model, we also provided insights on the duration of its effects, as this is key information to design valid pharmacological protocols in the perspective of a clinical application. The kinetics of reemergence of the former dysfunctions in SNpc DA neurons of the FXS mouse indicates that the ErbB-dependent effects are preserved for at least 4.5 h after intranigral drug injection and 7 h after systemic administration. Thus, a single treatment with the ErbB inhibitor produces lasting correction of the SNpc DA neuron dysfunctions underlying repetitive behaviors.

To date, there are no FDA-approved compounds for the treatment of repetitive stereotyped behaviors in patients with ASD and FXS. Current pharmacotherapy stands on off-label use of antipsychotics, which despite some positive effects, display adverse effects that undermine their therapeutic benefit^78^. Therefore, identifying novel therapeutic approaches to offset processes instrumental in repetitive behaviors remains an important mission. Besides gaining insight into the mechanisms that go awry in FXS, the present study demonstrate that targeting ErbB is an effective therapeutic strategy for aberrant repetitive behaviors. Beyond ASD and FXS, repetitive compulsive behaviors are hallmarks of different neuropsychiatric diseases, including obsessive-compulsive disorder and Tourette syndrome: rodent self-grooming is considered a translating phenotype of excoriation (compulsive skin picking), trichotillomania (compulsive hair pulling), dysmorphic disorder (obsessive cosmetic grooming) and tics, key features of these human diseases^79,80^. In this view, ErbB inhibition could be beneficial for the treatment of repetitive behaviors in different neuropsychiatric and neurodevelopmental diseases, besides ASD and FXS. Importantly, the therapeutic effectiveness also via systemic administration certifies the factual practicality of the ErbB inhibition-based approach, laying solid bases to propose ErbB targeting in clinical evaluations in ASD and FXS patients.

## Authors contributions

**SLD** performed mouse surgeries and intracerebral drug injections; performed and analyzed behavioral experiments. **ER** performed and analyzed western blot experiments. **MMC** performed and analyzed immunofluorescence experiments. **CB** supervised western blot experiments. **NBM** provided devices and resources. **AL** conceived the project, designed the experiments, performed and analyzed all electrophysiological recordings, dissected brain samples for western blot and immunofluorescence experiments, supervised the project, prepared the figures, and wrote the manuscript.

All authors discussed the results, commented on the manuscript and approved the final version.

## Acknowledgments

CB group work was supported by Telethon Foundation GGP20137, PRIN 20227JA8R3 MUR *Next Generation EU* and SNSF 310030 - 215706.

## Methods

### Mice

All procedures were conducted according to the guidelines on the ethical use of animals from the Council Directive of the European Communities (2010/63/EU) and were approved by the Italian Ministry of Health (authorization N°143-2020PR). *Fmr1* KO mice (C57/BL6J background) and C57/BL6J wild-type (WT) mice were obtained from Jackson Laboratories (USA) and then bred in our facility, housed in a temperature-(23 ± 1 ◦C) and humidity-controlled environment (45 %–60 % relative humidity) with a 12 h light/dark cycle. All experiments were performed in adolescent mice of 21-34 days of age.

### *In vivo* pharmacological treatments

#### Stereotaxic surgery and intracerebral drug injections

Male mice (26-28 days old) were anaesthetized with Zoletil 50 (Tiletamine HCl 4.1 mg/ml + Zolazepam HCl 4.1 mg/ml) and Rompun 20 (Xylazine 1.6 mg/ml) and mounted in a stereotaxic frame (David Kopf Instruments, Tujunga, CA) equipped with a mouse adapter. Mice were bilaterally implanted with a 26-gauge guide cannula positioned 1 mm above the SNpc using the following coordinates (from the brain surface): AP = − 3.4, ML = 1.2, DV = − 3.6 mm^81^. After 4-6 days of postoperative recovery, mice (30-32 days old) were bilaterally injected with PD158780 (10 µM) or it vehicle (Veh, 0.1 % DMSO in aCSF) through an injector cannula controlled by a microinfusion pump (total volume 0.6 μl/side at a continuous rate of 0.3 μl/min). The injector cannula was left for 2 min after the infusion to prevent backflow. Mice were returned to their home cage and after 30 min were subjected to procedures for electrophysiological experiments or behavioral tests.

#### Systemic drug injections

To evaluate the effects of systemic inhibition of ErbB signaling, mice (30-32 days old) were injected intraperitoneally (i.p.) with PD158780 (10 mg/Kg) or its vehicle (30% DMSO in NaCl 0.9 %) and after one hour were subjected to procedures for electrophysiological or behavioral analyses.

For intracerebral and systemic drug injections, PD158780 doses and times post-injection before starting electrophysiological or behavioral procedures have been chosen according to our previous *in vivo* evidence^33,82^ and other studies^83^.

#### Midbrain slice preparation

Acute midbrain slices containing the SNpc were used to perform electrophysiological recordings, immunofluorescence experiments, and SNpc microdissections for western blots. Midbrain slices were obtained following published procedures^84^. Briefly, mice (21-34 days old) were anesthetized with isoflurane and decapitated. The brain was quickly removed from the skull and a tissue block containing the midbrain was immersed in oxygenated aCSF at 8 °C, containing (in mM): NaCl 126, NaHCO_3_ 24, glucose 10, KCl 2.5, CaCl_2_ 2.4, NaH_2_PO_4_ 1.2 and MgCl_2_ 1.2 (95% O_2_–5% CO_2_, pH 7.3). Horizontal slices (250 μm) were cut using a vibratome (Leica VT1200S, Leica Microsystems, Wetzlar, Germany) and kept in aCSF at 33.0 ± 0.5°C for 30 min before being differently processed for distinct experiments.

#### Electrophysiological recordings

Patch-clamp recordings of SNpc DA neurons were performed with a setup equipped with an upright microscope (Nikon Eclipse FN1) and an infrared video-camera (CoolSNAP Photometrics). SNpc DA neurons were identified based on their morphology (fusiform, tightly packed, and medium to large-sized cell bodies), localization (in the medial SNpc, close to the medial terminal nucleus of the accessory optic tract, MT), and their typical electrophysiological features, as slow spontaneous firing (1–8 Hz) in cell-attached configuration, large action potential (>2ms) with a prominent afterhyperpolarization, or the presence of a prominent hyperpolarization-activated inward current (I_h_) in response to hyperpolarizing voltage steps^33,85^.

Cell-attached and whole-cell patch clamp recordings were performed with glass borosilicate pipettes (4–7 MΩ) pulled with a PP-83 Narishige puller and filled with a solution containing (in mM): 125 K-gluconate, 10 KCl, 10 HEPES, 2 MgCl_2_, 4 ATP-Mg_2_, 0.3 GTP-Na_3_, 0.75 EGTA, 0.1 CaCl_2_, 10 Phosphocreatine-Na_2_ (pH 7.3 with 2 M KOH). Recordings were made with a Multiclamp 700B amplifier (Molecular Devices, USA) using Clampex software (Molecular Devices, USA) and the Digidata 1440A A/D converter (Molecular Devices, USA) connected to a computer. The spontaneous firing activity was recorded in cell-attached configuration. In some experiments spontaneous firing was also recorded in whole-cell configuration (V_H_= -60 mV). Recordings were filtered at 1–4 kHz using the amplifier’s in-built low-pass filter and digitized at 10 kHz. The mean firing frequency and CV-IEIs were measured in at least 2-3 min of recordings. Membrane resistance (R_m_) and capacitance (C_m_) were measured with the “membrane test” protocol (Clampe× 10.3) consisting of a 5 mV hyperpolarizing step from -60 mV, at 33.3 Hz by averaging 100 consecutive measurements to calculate final values. R_m_ and C_m_ were measured within 1 min after membrane rupture. Membrane access resistance was repeatedly monitored and recordings in which it exceeded 25 MΩ were discarded. No liquid junction potential correction was applied.

To analyze neuronal excitability, a current-clamp protocol consisting of 2 s depolarizing current steps (+50/+250 pA, 50 pA step) was delivered to SNpc DA while maintaining their membrane potential at V_H_= -60 mV with current injection. The mean number of action potentials (APs) evoked by each current step was obtained by averaging results from three stimulation protocols.

To analyze E/I ratio in single nigral DA neurons, patch-pipettes were filled with an intracellular solution containing (in mM): Cs-methanesulfonate 115, CsCl 10, CaCl_2_ 0.45, HEPES 10, EGTA 1, QX-314 5, MgATP 4, NaGTP 0.3 (pH 7.3 with CsOH). Recordings of spontaneous excitatory postsynaptic currents (sEPSCs) were performed at V_H_= -70 mV, and spontaneous inhibitory postsynaptic currents (sIPSCs) were recorded in the same neuron at V_H_ = +10 mV and in presence of the glutamatergic AMPARs- and NMDARs antagonists, CNQX (10 µM) and APV (50 µM), respectively. sEPSCs and sIPSCs events were analyzed in 3 min of recordings.

The mGluR1/5 agonist (S)-3,5-dihydroxyphenylglycine (S)-DHPG (10 µM, 3 min) was perfused on midbrain slices to evoke mGluR1/5-mediated currents (I_DHPG_) in SNpc DA neurons. mGluR1- and mGluR5 contribution to I_DHPG_ was evaluated by measuring the residual I_DHPG_ following treatments with the mGluR1 antagonist CPCCOEt (100 µM, 15 min) or the mGluR5 antagonist MPEP (10 µM, 15 min), that unmask pure mGluR5- or mGluR1-mediated currents respectively^33^. I_DHPG_ amplitudes (V_H_= -60 mV) were calculated by measuring differences between baseline and peak.

The ErbB inhibitor PD15878 (IC_50_ for ErbB4 and ErbB2 is 52 nM and 49 nM, respectively) was used to assess effects induced by ErbB inhibition on nigral DA neurons. Concentration and duration of treatments were designed according to our previous evidence^33,34,82^. To evaluate the effects of acute ErbB inhibition in *ex vivo* patch-clamp recordings, PD158780 (10 µM in aCSF 0.1% DMSO) was perfused for 20-25 min on SNpc slices. To evaluate the effects of *in vivo* nigral ErbB inhibition, PD158780 (10 µM, 0.6 µL) or its vehicle (0.1 % DMSO in aCSF) were injected in SNpc, as described in *“Surgery and in vivo pharmacological treatments”*. After 30 min, mice were subjected to procedures for brain slicing and electrophysiology, and patch-clamp recordings were made up to 6.5 hours after intra-SNpc drugs injection.

In experiments to evaluate the effect of systemic ErbB inhibition on SNpc DA neurons’ functions, mice were i.p. treated with PD158780 (10 mg/Kg) or its vehicle (30% DMSO in NaCl 0.9 %). Brain slicing was performed after 1 h and electrophysiological recordings were made up to 7 h after systemic drug injections. (S)-DHPG, CPCCOEt and MPEP, CNQX and APV were purchased from Hellobio, while PD158780 was purchased from Abcam.

### Western blots

SNpc and VTA were dissected from midbrain slices of 25-day-old *Fmr1* KO and WT mice following published procedures^33,84^. Each sample was collected from three horizontal midbrain slices (250 μm) obtained from a single mouse. Midbrain slices, prepared as previously described, were kept in oxygenated aCSF at 33.0 ± 0.5 °C for 30 min (as for electrophysiological recordings). Then, SNpc and VTA dissection was performed in oxygenated aCSF at 33.0 ± 0.5 °C, and the collected samples were stored at -80 °C until further processing.

For western blot experiments, samples were homogenized with a tissue grinder dounce (WHEATON® Dounce Tissue Grinder 1 mL, VWR, Radnor, Pennsylvania, USA) in 150 mM NaCl (S3014 Sigma-Aldrich - Merck, Darmstadt, Germany), 50 mM Tris-HCl pH 7.5 (T1503 Sigma-Aldrich - Merck, Darmstadt, Germany), 1% Triton™ X-100 solution (93443 Sigma-Aldrich - Merck, Darmstadt, Germany), 1% sodium deoxycholate (D6750 Sigma-Aldrich - Merck, Darmstadt, Germany), 1 mM EDTA pH 8.0 (03690 Sigma-Aldrich - Merck, Darmstadt, Germany), 0.5 mM dithiothreitol (D0632 Sigma-Aldrich - Merck, Darmstadt, Germany), a protease inhibitor cocktail (P8340 Sigma-Aldrich - Merck, Darmstadt, Germany) and a phosSTOP-phosphatase inhibitor cocktail tablet (4906837001 Roche – Merck, Darmstadt, Germany). Brain region lysates were incubated for 10 minutes on ice and centrifuged for 10 minutes at maximum speed at 4°C. Protein concentration was measured using the Pierce™ BCA Protein Assay Kit (23225, Thermo Fisher Scientific, Waltham, Massachusetts, USA).

Proteins (30 μg) were separated on an 8% SDS-PAGE gel and blotted onto a PVDF membrane (GE10600029 Merck, Darmstadt, Germany). Membranes were blocked for 5 minutes with EveryBlot Blocking Buffer (12010020 Bio-Rad, Hercules, California, USA) following the manufacturer’s instructions. Membranes were incubated using the following primary antibodies: rabbit anti-ErbB4 (1:500, sc-283 Santa Cruz Biotechnology, Dallas, Texas, USA), rabbit anti-HER2/ErbB2 (1:500, 2165 Cell Signaling Technology, Danvers, Massachusetts, USA), mouse anti-Neuregulin-1/NRG1 (1:500, sc-393006 Santa Cruz Biotechnology, Dallas, Texas, USA), rabbit anti-mGluR1 (1:1000, 12551 Cell Signaling Technology, Danvers, Massachusetts, USA), mouse anti-Vinculin (1:2000, V9131 Sigma-Aldrich – Merck, Darmstadt, Germany) and mouse anti-β-Actin (1:5000, A5441 Sigma-Aldrich – Merck, Darmstadt, Germany). The following secondary antibodies were used: anti-rabbit IgG or anti-mouse IgG, HRP-linked Antibody (1:5000, 7074S and 7076S Cell Signaling Technology, Danvers, Massachusetts, USA). Proteins were revealed using an enhanced chemiluminescence kit (1705061 Bio-Rad, Hercules, California, USA) and the imaging system LAS-4000 mini (GE HealthCare Technologies Inc., Chicago, Illinois, USA). After incubation with antibodies against phospho-proteins, blots were stripped by incubation with Restore™ PLUS Western Blot Stripping Buffer (46430 Thermo Fisher Scientific, Waltham, Massachusetts, USA) according to manufacturer’s instructions. Each time after stripping, the absence of signal was confirmed before incubating the membrane with the antibody against the total protein. All phosphoproteins were normalized relative to the total protein on the same blot. Protein levels were normalized using the average of Coomassie blue staining, β-Actin and Vinculin signals on the membranes. Signal quantification was performed using ImageQuant TL software.

### Immunofluorescence and densitometric analysis

Midbrain slices were fixed in 4% paraformaldehyde in phosphate buffer (PB, 0.1 M, pH 7.4) overnight at 4 °C. Samples were rinsed three times in PB and immersed in 30% sucrose and 10% glycerol solution at 4 °C until sinking for cryoprotection. Slices were resliced into 30 μm-thick sections by a freezing microtome. Blocking was performed with 5% Normal Donkey serum (NDS) and 0.3% Triton X-100 (Sigma-Aldrich) in 0.1M PB solution. Free-floating sections were incubated with the primary antibodies for three nights at 4 °C in 5% NDS and 0.3% Triton X-100 in 0.1M PB solution. After three washes in PB, sections were incubated for 2 h with appropriate secondary antibodies. After three washes in PB, sections were mounted on slides, air-dried, and coverslipped with Fluoromount (Sigma Aldrich). The specificity of the immunofluorescence labeling was confirmed by omitting primary antibodies and using normal serum instead (negative controls). Confocal laser scanner microscopy (Zeiss, Oberkochen, Germany LSM800) was used to acquire images. All samples were acquired with the same laser settings and images were collected from 4-5 slices per animal (n=3). Quantitative analysis of TH/mGluR1-, TH/ErbB4-, or TH/ErbB2-positive neurons was performed with ImageJ software (NIH, USA). After background subtraction, the integrated density was measured for each individual cell. A range of 15-20 neurons was randomly analyzed for each slice. Data were presented as mean ± SEM and expressed as a percentage of control.

Primary antibodies: rabbit anti-mGluR1 (1:400, Cell Signalling; #12551), rabbit anti-ErbB4 (1:100, Santa Cruz; #283), rabbit anti-ErbB2 (1:100, Cell Signaling, #2165), and mouse anti-TH (1:1000, Millipore; #MAB318). Secondary antibodies: Alexa Fluor 488 donkey anti-mouse immunoglobulin G (1:200, Thermo Fisher Scientific; A-21202) and Alexa Fluor 555 donkey anti-rabbit immunoglobulin G (1:200, Thermo Fisher Scientific; A-31572).

### Behavior

All behavioural tests were conducted between 9:00 am and 16:00 pm after 1 hour of room acclimation.

#### Analyses of repetitive behaviors

A mouse was placed in a standard empty cage (24 cm x 13 cm x 20 cm) filled with fresh home-bedding, and video recorded for 15 min. First 5 min were considered as cage habituation, while the last 10 min were analysed for repetitive behaviors by an observer, evaluating time spent in self-grooming (stroking and licking face or body parts) and digging, and number of jumping.

#### Marble burying test

Marble burying test was conducted following published procedures^86^ with minor modifications. Briefly, a mouse was placed in a clean cage (40 cm x 18 cm x 26 cm) filled with fresh home-bedding (5 cm) and containing 20 marbles. The mouse was allowed to freely explore for 30 min. The number of marbles buried was calculated by considering buried a marble covered ≥ 50 % with bedding.

#### Open field

A mouse was introduced in the central part of the apparatus consisting of a circular open field (OF) (60 cm in diameter and 30 cm in height) and allowed to freely explore it for 30 min^82^. Locomotor activity was continuously recorded using a video camera placed above the arena; the total distance moved was automatically analysed using Ethovision XT 11 software (Noldus, The Netherlands). Then, the mouse was returned to the home cage. After each session, the apparatus was cleaned with 5% ethanol.

### Statistical analyses and reproducibility

Statistical analyses were conducted using OriginPro 2018b (OriginLab) or Prism9 (GraphPad). The normality of data sets was evaluated using Shapiro-Wilk or Kolmogorov-Smirnov test. For normally distributed data, statistical comparisons were performed by using two-tailed paired or unpaired Student’s *t*-test, two-way ANOVA, and repeated measures two-way ANOVA followed by post hoc tests, as appropriate. When normality was violated, data were analyzed with the non-parametric Mann–Whitney test or Kruskal-Wallis ANOVA followed by Kolmogorov-Smirnov test. Outlier values were evaluated with the Grubbs test and excluded from analyses. Data are represented as mean ± SEM and expressed as absolute or normalized values (as % of control). Statistical significance was set at P < 0.05, and indicated as * (P < 0.05), ** (P<0.01), and *** (P < 0.001), as appropriate. Sample sizes were not predetermined using statistical methods but based on previous experience with similar experimental settings. Experimenters were not blind to genotypes and drug treatments. Experiments were randomized whenever possible and replicated more times.

## Supplementary Figures

**Supplementary Fig. 1.**
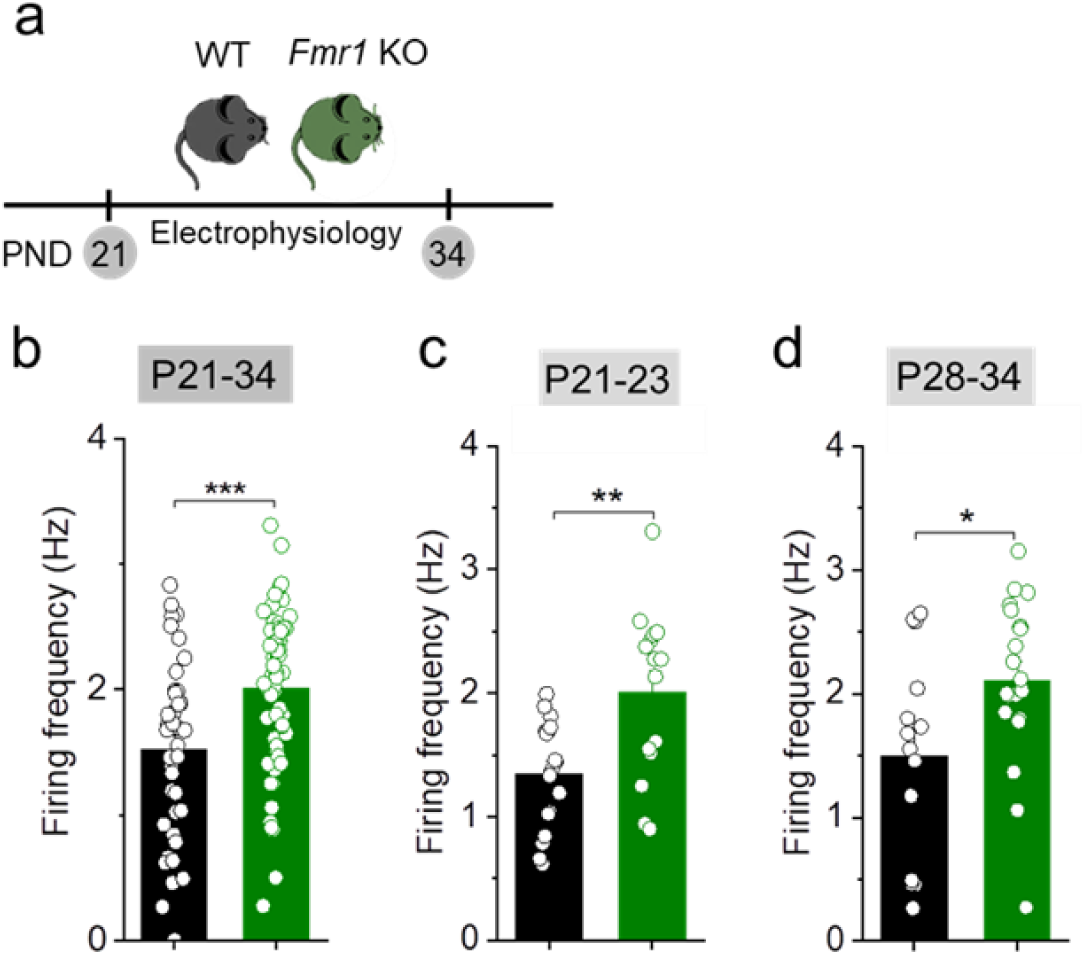
Nigral DA neuron hyperactivity is overt at the third postnatal week in the FXS mouse. **a)** Timeline of electrophysiological recordings of nigral DA neurons in adolescent *Fmr1* KO and WT mice. **b)** Plot of spontaneous firing frequency of nigral DA neurons from 21 to 34 day-old mice **(b)**. Quantification of firing rates at postnatal days 21-23 **(c)** or 28-34 **(d)** indicating that nigral DA neuron hyperactivity is an early feature of FXS evident at the third postnatal week in the FXS mouse. P21-34: WT (n = 41 cells/8 mice) and *Fmr1* KO (n = 59 cells/10 mice), P = 3.913 × 10-4, t = -3.671, two-tailed unpaired *t*-test. P21-23: WT (n = 18 cells/3 mice) and *Fmr1* KO (n = 15 cells/3 mice); P = 0.002, t = - 3.370, two-tailed unpaired *t*-test. P28-34: WT (n = 14 cells/3 mice) and *Fmr1* KO (n = 20 cells/3 mice); P = 0.0243, t = - 2.362, two-tailed unpaired *t*-test.

**Supplementary Fig. 2.**
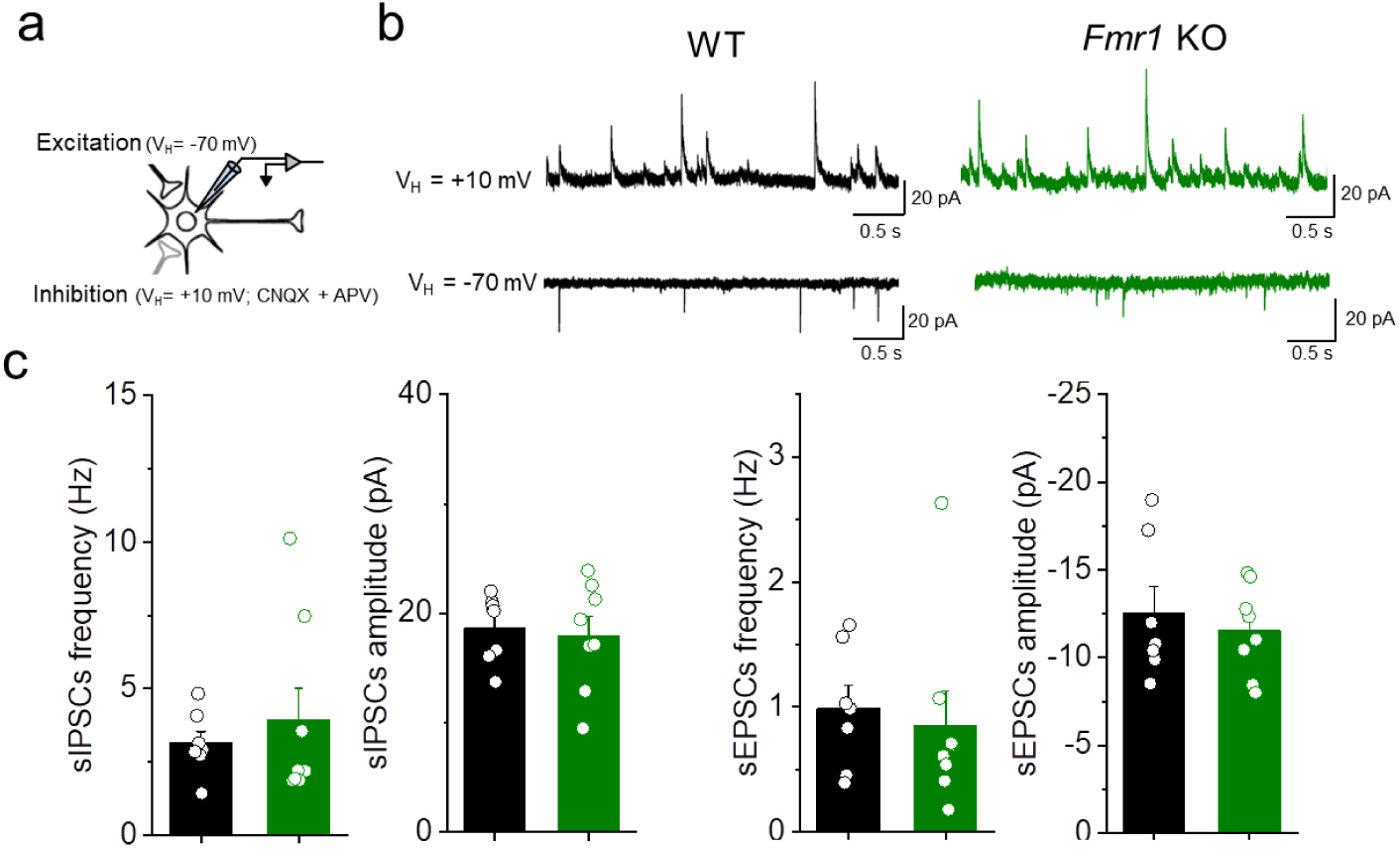
Normal excitation/inhibition (E/I) balance in nigral DA neurons of adolescent *Fmr1* KO mice. **a)** Scheme of experimental settings to record spontaneous excitatory postsynaptic currents (sEPSCs) and spontaneous inhibitory postsynaptic currents (sIPSCs) in the same nigral dopamine neuron. **b)** Example traces of sEPSCs and sIPSCs. **c)** Plots of mean frequencies and amplitudes of sIPSCs (left) and sEPSCs (right) showing no differences in excitatory and inhibitory inputs in SNpc DA neurons of WT (n = 8 cells/2 mice) and *Fmr1* KO mice (n = 9 cells/3 mice). sEPSCs frequency P = 0.463, U = 35, Mann-Whitney test; sEPSCs amplitude P = 0.955, U = 27, Mann-Whitney test; sIPSCs frequency P = 0.694, U = 32, Mann-Whitney test; sIPSCs amplitude P = 0.955, U = 27, Mann-Whitney test.

**Supplementary Fig. 3.**
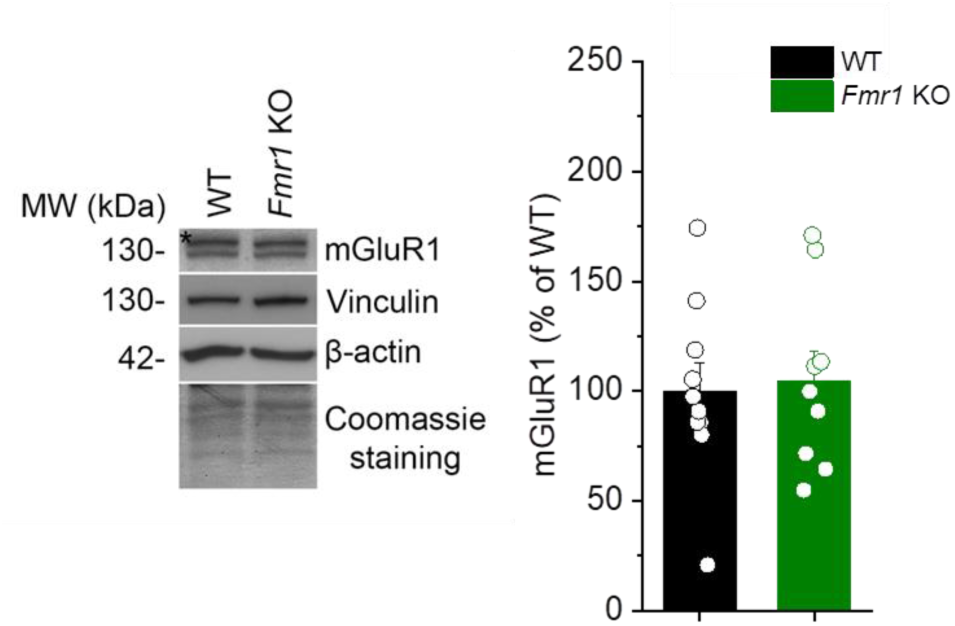
Normal mGluR1 levels in the ventral tegmental area (VTA) of adolescent *Fmr1* KO mice. Left, cropped representative western blots showing mGluR1 protein levels in total VTA from WT (n=10) and *Fmr1* KO mice (n=9). The molecular weight of each protein is indicated in kDa. Right, quantification of the western blots presented in the left panel. The graphs show the quantification of the signal intensity normalized over the WT condition (%) (P = 1, U = 45, Mann-Whitney test).

**Supplementary Fig. 4.**
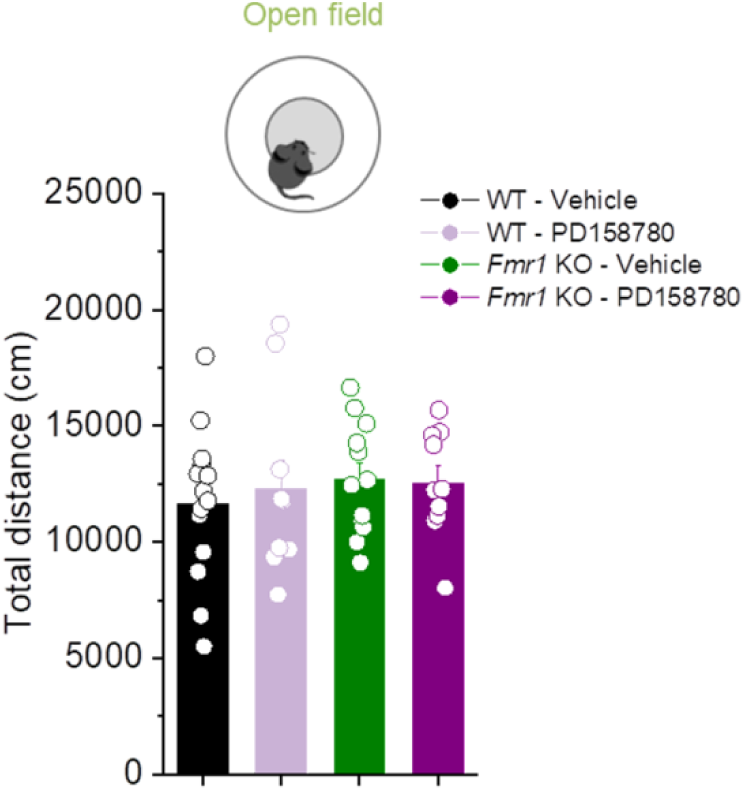
Effect of nigral ErbB inhibition on the locomotor activity of adolescent mice. Quantification of total distance moved in an open field by WT-Vehicle (n = 14), WT-PD158780 (n = 10), *Fmr1* KO-Vehicle (n=12), and *Fmr1* KO-PD158780 (n = 10). Two-way ANOVA. Genotype X drug interaction P = 0.645.

**Supplementary Fig. 5.**
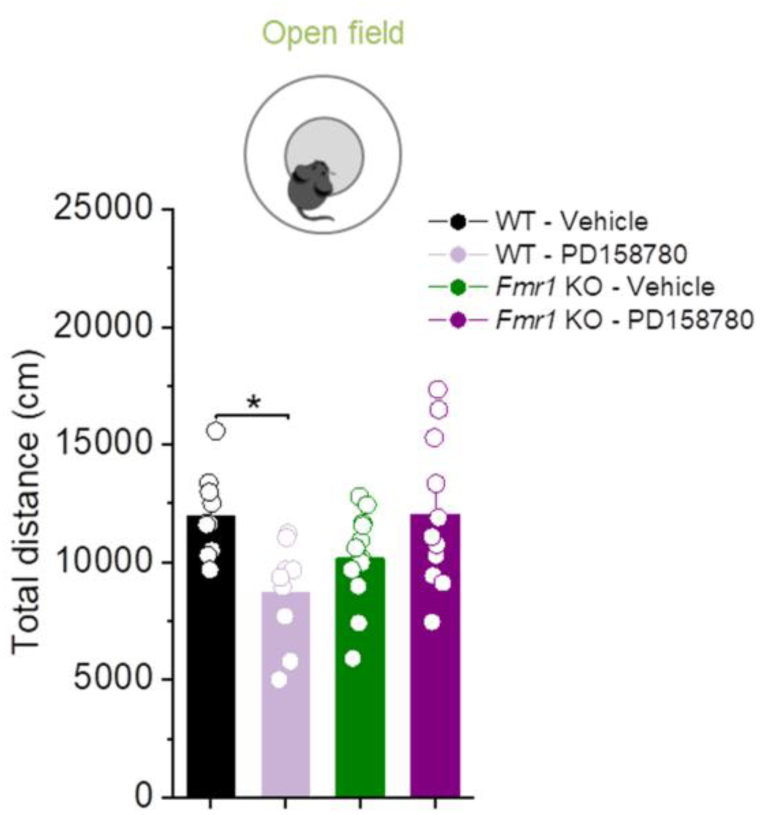
Effect of systemic ErbB inhibition on the locomotor activity of adolescent mice. Quantification of total distance moved in an open field by WT-vehicle (n = 10), WT-PD158780 (n = 9), *Fmr1* KO-vehicle (n = 13), and *Fmr1* KO-PD158780 (n = 11). Two-way ANOVA followed by Tukey’s test. ANOVA: genotype X drug interaction P = 9.060 × 10^-4^. P = 0.020 for WT-Vehicle vs WT-PD158780.

